# Splanchnic-spinal transmission of gut-initiated α-synucleinopathy

**DOI:** 10.64898/2026.10.19.760019

**Authors:** Byung-Kwon Jung, Min Kyung Song, Yeongseo Kim, Dae Ki Hong, Myounghun You, Eun Hee Ahn, Seong Su Kang, Tae-In Kam, Hyejin Park, Yoon-Seong Kim, Greg S. B. Suh

## Abstract

Gut-initiated α-synucleinopathy is proposed to spread from the gastrointestinal tract to the CNS, contributing to Parkinson’s disease (PD), but whether routes beyond the vagus nerve are involved remains unclear. Using complementary intracolonic rotenone and gastroduodenal α-synuclein PFF models, we combined pathway-specific surgical interruption with longitudinal behavioral analysis and anatomical mapping of pathological α-synuclein. In the intracolonic rotenone model, interruption of the prevertebral ganglia/splanchnic–spinal axis more strongly attenuated behavioral deficits and α-synuclein pathology than vagotomy. In the gastroduodenal PFF model, both vagal and splanchnic–spinal pathways contributed to pathology propagation. Early after PFF inoculation, pathological α-synuclein was detected in peripheral neural structures associated with both routes. Consistent with these findings, spinal pathology was evident in human PD. Together, these results identify the splanchnic–spinal pathway as a key component of body-first Parkinson’s disease, broadening current models of peripheral-to-central disease progression and informing efforts to target early peripheral disease progression.

## Introduction

Parkinson’s disease (PD) is a progressive neurodegenerative disorder characterized by degeneration of nigrostriatal dopaminergic neurons and the accumulation of misfolded α-synuclein (α-Syn) in Lewy bodies and Lewy neurites. The identification of α-Syn as a major component of Lewy pathology, together with experimental evidence that fibrillar α-Syn can seed endogenous α-Syn aggregation and propagate pathology through interconnected neuronal populations, has supported the idea that neural transmission of pathological α-Syn may contribute to disease progression **(Spillantini et al., 1997; Volpicelli-Daley et al., 2011; Luk et al., 2012)**. This concept converged with the staging scheme proposed by Braak and colleagues, in which α-Syn pathology was observed in the lower brainstem and olfactory structures before progressively involving more rostral regions, and with subsequent observations of α-Syn pathology in the enteric nervous system (ENS) **(Braak et al., 2003, 2006)**. These findings led to the hypothesis that, in at least a subset of patients, α-synucleinopathy may originate in peripheral tissues, including the gastrointestinal tract, and subsequently spread toward the central nervous system (CNS). Although this sequence does not account for all forms of PD, converging clinical, imaging, and neuropathological evidence supports a body-first trajectory, in which peripheral α-synucleinopathy may precede CNS involvement in a substantial proportion of patients **(Horsager and Borghammer, 2024)**.

Among the neural pathways connecting the gut to the CNS, the vagus nerve has become the principal candidate for mediating gut-to-brain α-Syn transmission. Its prominence reflects a convergence of anatomical, pathological, epidemiological, and experimental evidence. Vagal afferents provide a major sensory pathway conveying gastrointestinal information to the hindbrain **(Berthoud et al., 2004)**. Notably, the dorsal motor nucleus of the vagus (DMV), a key parasympathetic output nucleus of the vagovagal circuit, is among the brainstem regions affected early in the Braak staging scheme **(Braak et al., 2003)**.

Importantly, registry studies in Denmark and Sweden reported a lower subsequent risk of PD after truncal vagotomy, particularly with longer follow-up **(Svensson et al., 2015; Liu et al., 2017)**. This pattern supports a role for vagal continuity in disease development, although the observational nature of these studies precludes causal inference. Moreover, because registry data cannot establish the anatomical exclusivity of individual operations, potential effects on adjacent autonomic structures or non-vagal gut-connected neural elements, including components of the splanchnic–spinal circuitry, cannot be fully excluded. Nevertheless, experimental studies have supported a role for the vagal pathway in α-Syn transfer. α-Syn introduced into the gut can access vagally connected structures **(Holmqvist et al., 2014)**, and gastroduodenal α-Syn PFF inoculation produces progressive CNS α-synucleinopathy that is strongly suppressed by truncal vagotomy **(Kim et al., 2019)**. These results have established the vagus nerve as the dominant neural route in current models of gut-originating α-synucleinopathy.

The vagus nerve, however, constitutes only one component of the extrinsic neural circuitry linking the gastrointestinal tract to the CNS. Gut-derived sensory information is conveyed through anatomically distinct vagal and spinal pathways: vagal afferent neurons reside primarily in the nodose ganglia and project centrally to the hindbrain, whereas spinal afferent neurons reside in dorsal root ganglia and enter the CNS through the spinal cord **(Sharkey and Mawe, 2023)**. Many gut-projecting spinal afferents travel through splanchnic pathways, often alongside sympathetic efferent fibers, while remaining neuroanatomically distinct sensory populations **(Berthoud et al., 2004; Wolfson, 2025)**. In parallel, sympathetic output to the gastrointestinal tract is organized largely through thoracolumbar preganglionic neurons and postganglionic neurons within prevertebral ganglia, including the celiac and superior mesenteric ganglia **(Sharkey and Mawe, 2023)**. These vagal and spinal pathways convey partially overlapping but functionally distinct information about the gastrointestinal environment, with spinal afferents contributing prominently to mechanosensory, inflammatory, chemical, and nociceptive signaling **(Berthoud et al., 2004; Wolfson, 2025)**. Thus, gut–brain communication is inherently organized through parallel extrinsic neural routes rather than through the vagus nerve alone. Notably, α-Syn pathology in human Lewy body disorders is likewise not confined to vagally connected structures: phosphorylated α-Syn has been observed in the spinal cord and paraspinal sympathetic ganglia, including autonomic spinal regions **(Beach et al., 2010)**. Experimental rotenone studies have further suggested that enteric α-Syn can be taken up by sympathetic neurites **(Pan-Montojo et al., 2012)**. Yet whether a gut-connected splanchnic– spinal pathway serves as an ascending route for pathological α-Syn to reach the spinal cord and contribute to subsequent CNS pathology, and how much this route contributes relative to the established vagal pathway, remains unresolved.

Here, we investigated the neural routes through which gut-initiated α-synucleinopathy reaches the CNS, with particular emphasis on the contribution of the splanchnic–spinal pathway in addition to the established vagal route. Using complementary intracolonic rotenone and gastroduodenal α-Syn PFF models, we combined surgical interruption of distinct gut–brain neural pathways with longitudinal behavioral analyses and anatomical mapping of α-Syn pathology across peripheral nerves, ganglia, the spinal cord, and the brain. We further examined human postmortem tissue to assess the relevance of spinal α-Syn pathology to PD. Together, our findings identify the splanchnic–spinal pathway as a major component of gut-to-brain α-Syn pathology and support a model in which body-first PD can engage parallel neural routes to the CNS.

## Results

### Intracolonic Rotenone Administration Drives Progressive Gut-to-Brain α-Synuclein pathology

Although gut-initiated α-synucleinopathy can propagate to the brain via the vagus nerve (parasympathetic system), splanchnic-spinal nerves also densely innervate the gastrointestinal tract, representing another physical pathway connecting the gut and the central nervous system. To investigate which neural pathways mediate the transmission of gut-initiated α-synucleinopathy, we established a novel gut-initiated mouse model using intracolonic administration of rotenone, a chemical inducer of intestinal synucleinopathy. We deliberately chose the intracolonic route over conventional oral administration for two reasons: 1) To eliminate systemic toxicity, as systemic rotenone exposure often leads to widespread non-specific pathological confounding factors, and 2) To target the posterior gut (cecum and colon), where gut microbiota—whose alterations are linked to α-synuclein pathology and Parkinson’s disease—are primarily localized, thereby allowing precise local exposure without chronic whole-gut disruption. To track the spatiotemporal pathology, mice received chronic intracolonic injections of rotenone for 6 weeks, and intestinal alterations were systematically analyzed at both early and late time points **(Figure 1A and Figure S1A and S1B)**.

**Figure 1.**
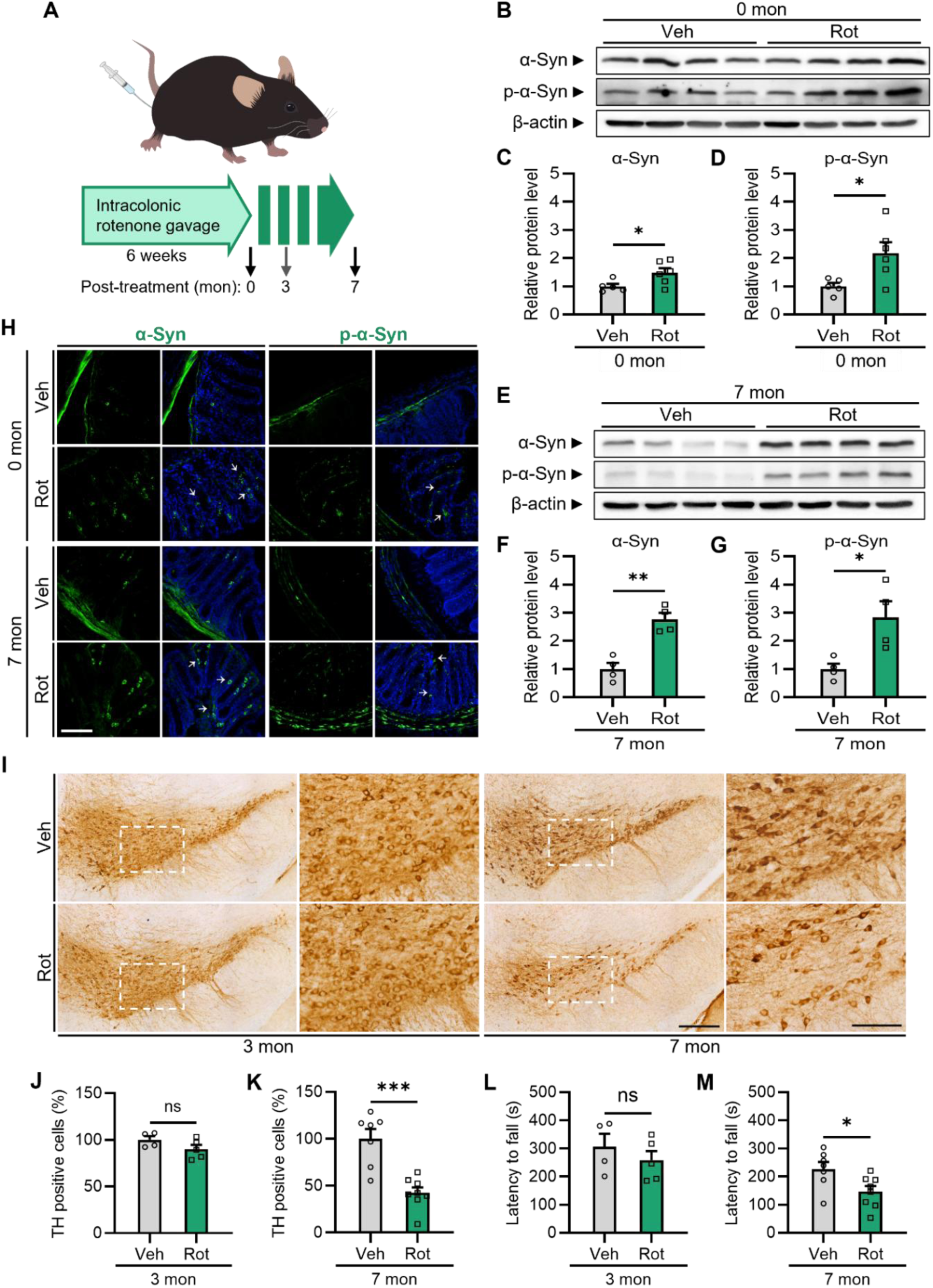
Intracolonic administration of rotenone induces early α-Syn pathology in the colon and late-onset dopaminergic neurodegeneration in the brain. **(A)** Schematic illustration of the experimental procedure for intracolonic rotenone administration and the corresponding timeline. 0 mon indicates the time point immediately following completion of the 6-week rotenone treatment; 3 mon and 7 mon represent 3 months and 7 months post-treatment, respectively. **(B)** Representative immunoblots showing colonic protein levels of α-Syn, phosphorylated α-Syn (p-α-Syn, Ser129), and β-actin at 0 mon. **(C and D)** Quantification of colonic α-Syn **(C)** and p-α-Syn **(D)** protein levels normalized to β-actin at 0 mon (n = 4 per group). **(E)** Representative immunoblots showing colonic protein levels of α-Syn, p-α-Syn, and β-actin at 7 mon post-treatment. **(F and G)** Quantification of colonic α-Syn **(F)** and p-α-Syn **(G)** protein levels normalized to β-actin at 7 mon post-treatment (n = 4 per group). **(H)** Representative immunofluorescence images of the colon stained for α-Syn (green, left) or p-α-Syn (green, right), with DAPI counterstaining (blue), at 0 mon and 7 mon post-treatment. Arrows indicate α-Syn-or p-α-Syn-positive cells. Scale bar, 50 μm. **(I)** Representative immunohistochemical images of tyrosine hydroxylase (TH)-positive dopaminergic neurons in the substantia nigra pars compacta (SNpc) at 3 mon and 7 mon post-treatment. White dashed boxes indicate regions shown at higher magnification on the right. Scale bars, 300 μm (main images) and 100 μm (insets). **(J and K)** Quantification of relative TH-positive cell counts in the SNpc at 3 mon **(J)** and 7 mon (K) post-treatment (n = 4–8 per group). **(L and M)** Rotarod performance of vehicle-or rotenone-treated mice at 3 mon (L) and 7 mon (M) post-treatment (n = 4–8 mice per group). Data are presented as mean ± SEM. Statistical significance was determined using an unpaired two-tailed Student’s t-test; \**p* < 0.05, \*\**p* < 0.01, \*\*\**p* < 0.001; ns, not significant. Veh, vehicle; Rot, rotenone.

Chronic intracolonic administration of rotenone did not induce significant changes in body weight **(Figure S1C)**. However, it upregulated both total α-synuclein and phosphorylated α-synuclein at Serine 129 (pS129 α-syn), a key pathological marker of synucleinopathy, in the colon at the protein level. Specifically, immunoblot analysis revealed significant increases in α-synuclein and pS129 α-syn protein levels in the colon at both the early **(Figure 1B–D)** and late stages **(Figure 1E–G)**. Notably, while α-synuclein mRNA expression remained unaltered immediately following treatment **(Figure S1D)**, its transcriptional level was significantly elevated at 7 months post-treatment **(Figure S1E)**, indicating a delayed transcriptional upregulation accompanied by sustained pathological protein accumulation over time. Additionally, histological analysis revealed a marked increase in pS129 α-syn-positive cells within the colonic mucosa, accompanied by elevated signals of proteinase K-resistant α-synuclein aggregates at the tissue level **(Figure 1H and Figure S1F)**. To assess early mucosal responses, we evaluated intestinal barrier integrity and inflammation. Protein levels of tight junction components, Claudin-1 and Occludin, remained largely unchanged at the early time point **(Figures S2A–S2C)**, but significantly altered at 7 months post-treatment **(Figures S2A, S2D, and S2E)**. Notably, fecal levels of Lipocalin-2 (Lcn2), a marker of intestinal inflammation, were significantly elevated immediately following treatment **(Figure S2F)**, followed by a marked increase in *IL6* mRNA expression at 7 months **(Figure S2G)**. These results indicate that chronic intracolonic exposure to rotenone triggers an early inflammatory response in the gut. Likely associated with this response, reduced gut motility emerges at an early stage (3 months), leading to constipation-like phenotypes that persist into the late stage **(Figures S1G–S1N)**.

In contrast to the early-onset intestinal alterations, brain pathology—characterized by α-synuclein aggregation and dopaminergic loss—was absent immediately, becoming evident predominantly at the late stage, as represented by the 7-month post-exposure timepoint **(Figures 1I–1K)**. Behaviorally consistent with these findings, motor deficits were absent during the early stage, whereas a noticeable decline in motor performance was observed at the late stage **(Figures 1L and 1M)**. Altogether, these findings demonstrate that chronic intracolonic administration of rotenone establishes a progressive model of gut-initiated α-synucleinopathy, linking early-onset gut alterations to the late-onset neuropathology.

### Surgical Transection of Gut–Brain Neural Pathways Attenuates Rotenone-Induced Behavioral Impairments

To assess the individual contribution of each gut-to-brain neural connection to intracolonic rotenone-induced brain pathology, we performed surgical transection of either the vagal or spinal pathway innervating the gut. For effective and permanent blockade of these neural pathways, subdiaphragmatic truncal vagotomy (SDVx) was performed by bilaterally transecting the vagal trunks above the hepatic branch along the esophagus **(Figure 2A)**, while celiac–superior mesenteric ganglionectomy (CGx) was executed by resecting the prevertebral CSMG complex with additional transection of adjacent sympathetic branches **(Figure 2B)**. Following these surgical procedures, post-hoc validation was performed to confirm successful neural blockade, and rule out functional regeneration and weight loss **(Figures S3A-S3C)**. After a recovery period, the impact of these denervations on rotenone-induced pathology was evaluated through a battery of behavioral assessments **(Figure 2C-2H)**.

**Figure 2.**
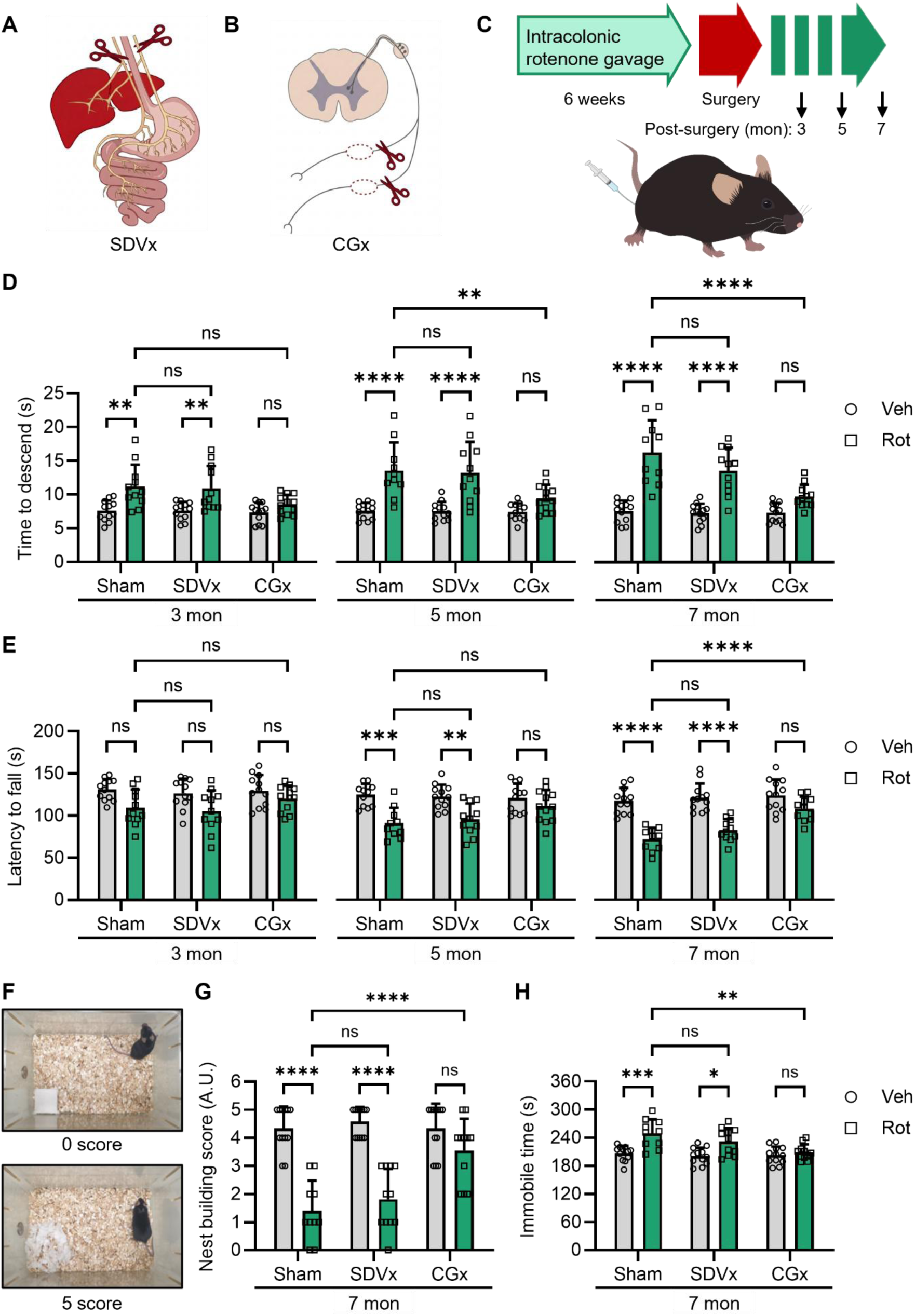
Celiac-superior mesenteric ganglionectomy ameliorates Parkinson’s disease-related pathologic symptoms induced by intracolonic rotenone administration. **(A)** Schematic illustration of subdiaphragmatic truncal vagotomy (SDVx). **(B)** Schematic illustration of celiac-superior mesenteric ganglionectomy (CGx). **(C)** Experimental timeline showing intracolonic rotenone administration, surgical denervation procedures (Sham, SDVx, or CGx), and behavioral assessments at 3, 5, and 7 mon post-surgery. **(D)** Pole-test performance, measured as total time to descend, in vehicle (Veh)-and rotenone (Rot)-treated mice at 3, 5, and 7 mon post-treatment following Sham, SDVx, or CGx surgery (n = 10–12 mice per group). **(E)** Rotarod performance, measured as latency to fall, in Veh-and Rot-treated mice at 3, 5, and 7 mon post-treatment following Sham, SDVx, or CGx surgery (n = 10–12 mice per group). **(F)** Representative photographs of nest-building behavior showing an intact Nestlet corresponding to a score of 0 (top) and a fully formed nest corresponding to a score of 5 (bottom). **(G)** Quantification of nest-building scores in Veh-and Rot-treated mice at 7 mon post-treatment following Sham, SDVx, or CGx surgery (n = 10–12 mice per group). **(H)** Tail-suspension immobility time in Veh-and Rot-treated mice at 7 mon post-treatment following Sham, SDVx, or CGx surgery (n = 10–12 mice per group). Data are presented as mean ± SEM. Statistical significance was determined by two-way ANOVA followed by Tukey’s or Sidak’s multiple-comparisons test, as appropriate; \**p* < 0.05, \*\**p* < 0.01, \*\*\**p* < 0.001; ns, not significant. Veh, vehicle; Rot, rotenone.

Longitudinal evaluation using the pole test demonstrated that intracolonic rotenone administration induced a progressive motor deficit from 3 to 7 months post-administration compared with vehicle controls **(Figure 2D)**. Notably, the rotenone-induced motor deficit persisted in the vagotomy (SDVx) group; however, this behavioral impairment was markedly attenuated in CGx-treated mice. These behavioral outcomes were further supported by the rotarod test: while mice receiving SDVx showed a slight, non-significant amelioration, those in the CGx group maintained motor performance at levels comparable to vehicle controls despite rotenone exposure **(Figure 2E)**. Extended evaluation up to 9 months post-administration revealed that SDVx exhibited a modest rescue effect in both the pole and rotarod tests; however, mice in the CGx group continuously maintained motor performance at levels comparable to vehicle controls **(Figures S3D and S3E)**. Next, we assessed sensorimotor coordination using the nest-building assay. The rotenone-induced decline in nest-building performance was not affected by SDVx but was selectively attenuated by CGx **(Figures 2F and 2G)**. Similarly, the rotenone-induced depression-like behavior assessed by the tail-suspension test was selectively abolished by CGx, but not by SDVx **(Figure 2H)**.

To determine whether these protective effects were uniquely attributable to our localized intracolonic model, we evaluated the impact of surgical denervation on behavioral outcomes in mice subjected to conventional oral rotenone administration. Unlike chronic intracolonic administration, mice subjected to chronic oral administration exhibited immediate motor deficits following treatment **(Figures S3F and S3G)**. Because surgical interventions (Sham, SDVx, and CGx) require a recovery period and their immediate interactions with oral rotenone exposure were uncertain, we performed surgical denervations immediately following chronic oral rotenone administration. After an adequate recovery phase, motor performance was evaluated at 3 months post-administration using the pole and rotarod tests. Both SDVx and CGx modestly attenuated oral rotenone-induced motor deficits; however, motor function in both surgical groups remained significantly impaired compared with that in vehicle controls **(Figures S3H and S3I)**.

Nevertheless, whether the acute motor deficits ensued immediately after oral administration stem from authentic gut-to-brain α-synuclein propagation remains unclear. Furthermore, it is difficult to determine whether the mild protective effects of denervation observed at 3 months result from blocked pathological transmission or from off-target neurophysiological alterations secondary to surgery. Taken together, behavioral evaluation in our intracolonic rotenone model demonstrates that, in addition to the classical vagal route, spinal nerves innervating the gut also contribute to progressive Parkinson’s disease-like pathology, highlighting the spinal pathway as an alternative route for pathological transmission.

### Celiac Sympathetic Denervation Attenuates Pathological α-Synuclein Propagation along the Spinal-to-Brain Axis

Having shown that denervation of the splanchnic nerve attenuated the gut-initiated PD pathogenesis, we next evaluated the histological mechanism underlying the observed behavioral protection by examining whether surgical denervation affected the accumulation and propagation of phosphorylated α-synuclein (pS129-α-Syn) along the spinal cord and hindbrain. Immunohistochemical analysis of the cervical spinal cord revealed a robust increase in pS129-α-Syn immunoreactivity in both the dorsal and ventral horns following intracolonic rotenone administration **(Figure 3A)**. Consistent with the anatomical distinctiveness of these neural pathways, SDVx failed to prevent rotenone-induced pS129-α-Syn accumulation in the spinal cord; by contrast, CGx almost completely blocked the accumulation of pathological pS129-α-Syn in both dorsal and ventral regions **(Figures 3A–3C)**.

**Figure 3.**
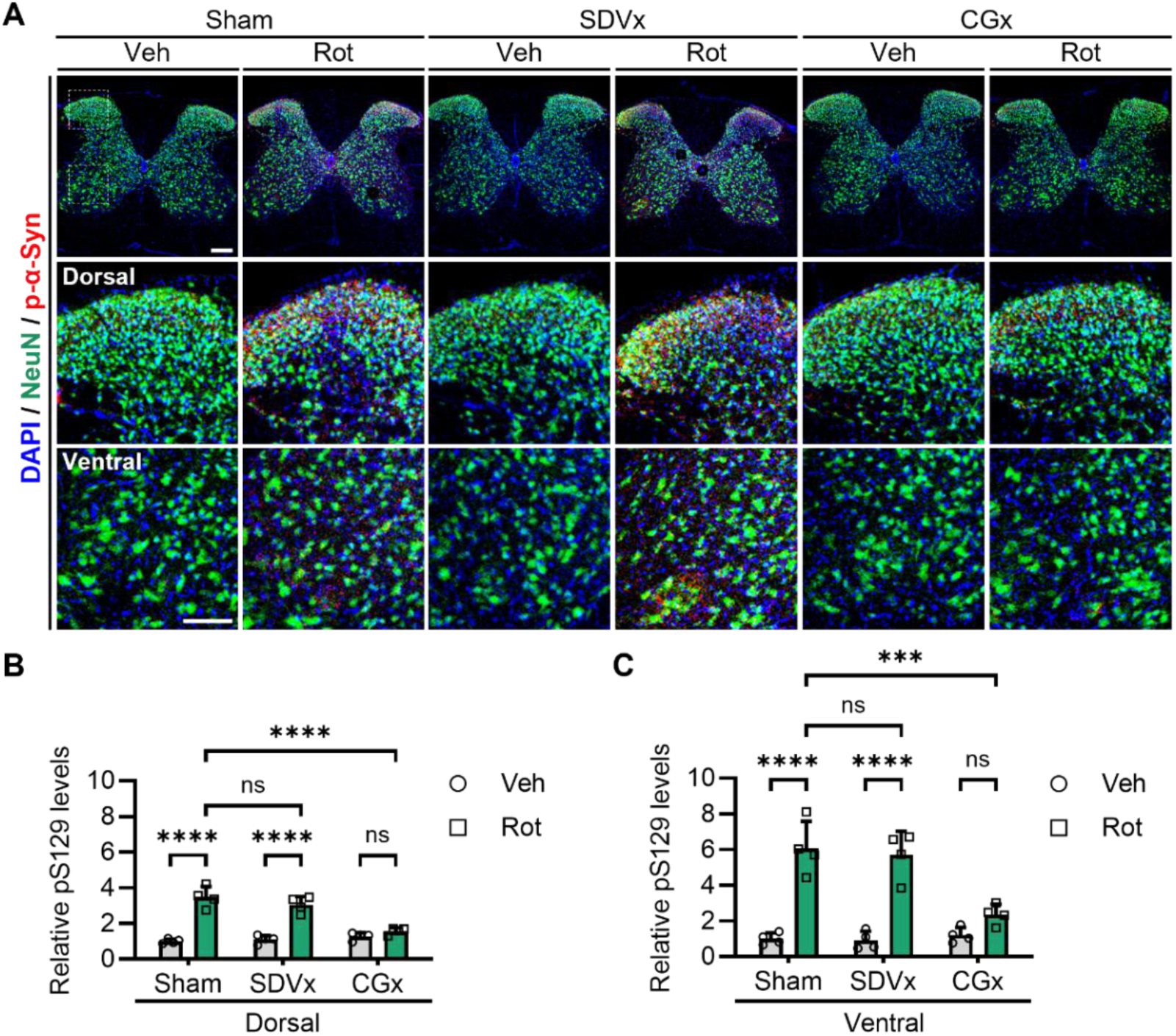
Celiac-superior mesenteric ganglionectomy, but not vagotomy, prevents pathological α-synuclein accumulation along the spinal-to-brain axis in rotenone-treated mice. **(A)** Representative immunofluorescent images of coronal cervical spinal cord sections collected at 7 months post-administration, stained for NeuN (green), p-α-Syn (red), and DAPI (blue). High-magnification images of the dorsal horn (middle) and ventral horn (bottom) correspond to the dashed boxes in the top panel. Scale bars, 200 *μ*m (top) and 50 *μ*m (magnified). **(B and C)** Relative quantification of p-α-Syn levels in the dorsal horn (B) and ventral horn (C) of the cervical spinal cord (*n* = 4 per group). Data are presented as mean ± SEM. Statistical significance was determined by one-way ANOVA followed by Tukey’s multiple comparisons test; \**p* < 0.05, \*\**p* < 0.01, \*\*\**p* < 0.001; ns, not significant. Veh, vehicle; Rot, rotenone; SDVx, subdiaphragmatic truncal vagotomy; CGx, celiac–superior mesenteric ganglionectomy.

### Exogenous α-Syn Preformed Fibril Induces α-Synucleinopathy from the Gut to the Brain through both Vagal and Spinal pathways

While rotenone administration models α-synucleinopathy, its potential to induce widespread cytotoxicity—such as global mitochondrial dysfunction and elevated oxidative stress independent of direct α-synuclein pathology—cannot be entirely ruled out. Thus, to further extend our hypothesis, we utilized exogenous α-Syn preformed fibrils (PFF). While the established gut-first model via microinjection of α-Syn PFF into the myenteric plexus of the upper small intestine provides an α-synuclein-dependent proteinopathy framework and has been shown to propagate through the vagus nerve **(Kim et al., 2019)**, its potential propagation via the spinal/sympathetic axis remains unexplored. To investigate that, we injected α-Syn preformed fibrils (PFF) into the mouse upper duodenum stomach wall and performed surgical denervation of either the vagal or sympathetic pathways **(Figure 4A)**. We also evaluated the efficiency and physiological impact of surgical denervations in this work: SDVx led to a noticeable increase in gross stomach size and wet weight, characteristic of impaired gastric motility following vagal nerve transection, whereas CGx did not alter stomach morphology compared to PBS controls **(Figure S4A and S4B)**.

**Figure 4.**
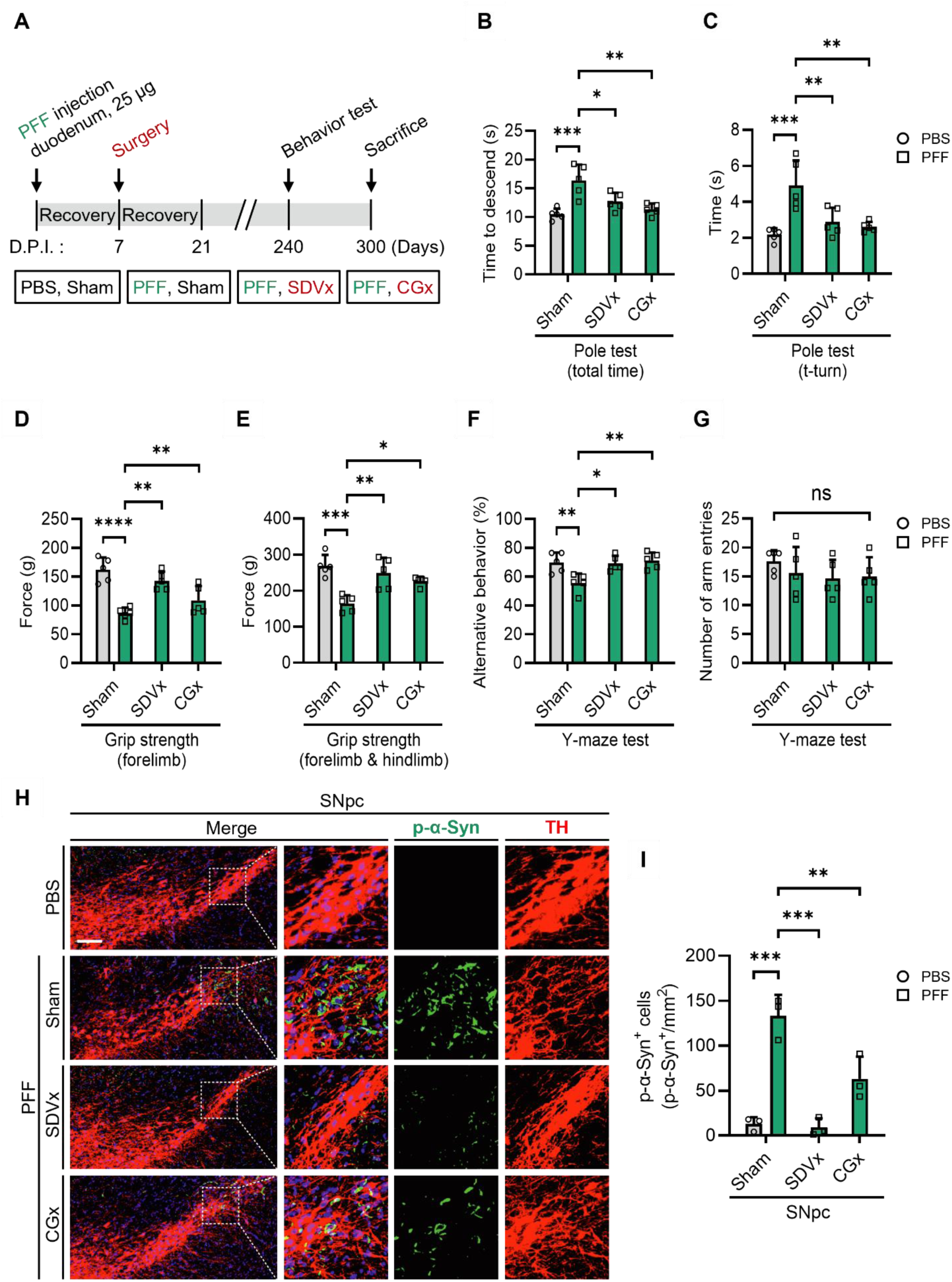
Celiac-superior mesenteric ganglionectomy or vagotomy prevents α-synuclein preformed fibrils-induced Parkinson’s disease-related pathological symptoms. **(A)** Experimental timeline illustrating upper gastrointestinal administration of α-synuclein preformed fibrils (α-Syn PFF), surgical denervation procedures (Sham, SDVx, or CGx), and sequential behavioral and histological evaluations at 8 or 10 months post-injection. **(B and C)** Behavioral assessment using the pole test in PBS-or PFF-injected mice subjected to Sham, SDVx, or CGx surgery at 8 months post-injection (*n* = 5 mice per group). **(D and E)** Grip strength measurements of the forelimbs (D) and combined forelimbs and hindlimbs (E) in PBS-or PFF-injected mice subjected to Sham, SDVx, or CGx surgery at 8 months post-injection (*n* = 5 mice per group). **(F and G)** Behavioral assessment of short-term or working memory using the Y-maze test in PBS-or PFF-injected mice subjected to Sham, SDVx, or CGx surgery at 8 months post-injection (*n* = 5 mice per group). Quantification showing the percentage of spontaneous alternations (F) and total number of arm entries (G) during the Y-maze test. **(H)** Representative immunofluorescent images of tyrosine hydroxylase (TH, red)-positive dopaminergic neurons and p-α-Syn (green) aggregates with DAPI counterstain (blue) in the substantia nigra pars compacta (SNpc) at 10 months post-injection. White dashed boxes indicate magnified regions shown on the right. Scale bars, 200 *μ*m. **(I)** Density of p-α-Syn-positive cells (cells/mm^2^) in the SNpc evaluated at 8 months post-injection (*n* = 3 per group). Data are presented as mean ± SEM. Statistical significance was determined by one-way ANOVA followed by Tukey’s multiple comparisons test; \**p* < 0.05, \*\**p* < 0.01, \*\*\**p* < 0.001; ns, not significant. PBS, phosphate-buffered saline; PFF, preformed fibrils; SDVx, subdiaphragmatic truncal vagotomy; CGx, celiac–superior mesenteric ganglionectomy.

Having confirmed successful surgical denervations, we next assessed the functional consequences on motor behavior. Intragastric injection of PFF induced marked motor deficits and behavioral impairments **(Figure 4B–4G)**, consistent with the previous report **(Kim et al., 2019)**. Remarkably, surgical blockade of either the vagal or sympathetic route (SDVx or CGx) partially rescued these peripheral PFF-induced motor deficits, indicating that both neural pathways actively contribute to functional impairments **(Figure 4B–4G)**.

To determine whether these functional changes were accompanied by changes in the underlying anatomical substrate, we examined whether vagal and spinal denervations differentially block the retrograde transport of pathological α-Syn to the brainstem. Gastric injection of PFF robustly induced phosphorylated α-Syn (p-α-Syn) accumulation in the dorsal motor nucleus of the vagus nerve (DMV). While CGx led to a modest reduction in p-α-Syn levels within the DMV, SDVx produced a substantially greater attenuation of p-α-Syn accumulation in cholinergic neurons (ChAT⁺) **(Figure S4C and S4D)**. These findings confirm that vagal efferents/afferents serve as the primary and predominant transmission route of pathological α-Syn to the parasympathetic brainstem center. Furthermore, we found that gastric PFF administration led to a noticeable loss of tyrosine hydroxylase-positive (TH⁺) neurons in the LC. Notably, SDVx almost completely prevented this PFF-induced loss of TH⁺ neurons, restoring levels comparable to PBS-treated controls **(Figure S4E and S4F)**. On the other hand, CGx exhibited only modest amelioration of TH⁺ neuronal loss, indicating that vagal integrity plays a crucial role in preventing LC neurodegeneration in this gastric PFF model **(Figure S4E and S4F)**.

We next sought to determine whether this gut-derived pathology further ascends to the midbrain dopaminergic system to drive neurodegeneration. PFF injection into the duodenum caused prominent p-α-Syn accumulation in tyrosine hydroxylase-positive (TH⁺) neurons of the substantia nigra pars compacta (SNpc) **(Figure 4H and 4I)**. Notably, both SDVx and CGx significantly reduced p-α-Syn pathology in the SNpc, suggesting that both vagal and spinal pathways independently contribute to the ascending spread of α-synucleinopathy toward midbrain dopaminergic regions **(Figure 4H and 4I)**.

To directly determine whether gut-injected α-Syn PFF propagates through the spinal cord, we examined p-α-Syn accumulation in both thoracic and cervical levels of the spinal cord. PFF administration markedly elevated p-α-Syn signal intensity in both the dorsal and ventral horns of thoracic and cervical spinal cord sections **(Figure 5A–5F)**. Crucially, while SDVx did not block this spinal cord pathology, CGx significantly attenuated p-α-Syn accumulation in both thoracic and cervical segments, demonstrating that the spinal/sympathetic axis specifically mediates retrograde transmission of pathological α-Syn to the spinal cord. Taken together, these results demonstrate that exogenous α-Syn PFF injected into the gut propagates to the central nervous system not only through the vagal/parasympathetic nerve to the brainstem, but also via the spinal/sympathetic pathway through the spinal cord, with both conduits converging on the midbrain and driving widespread pathological outcomes.

**Figure 5.**
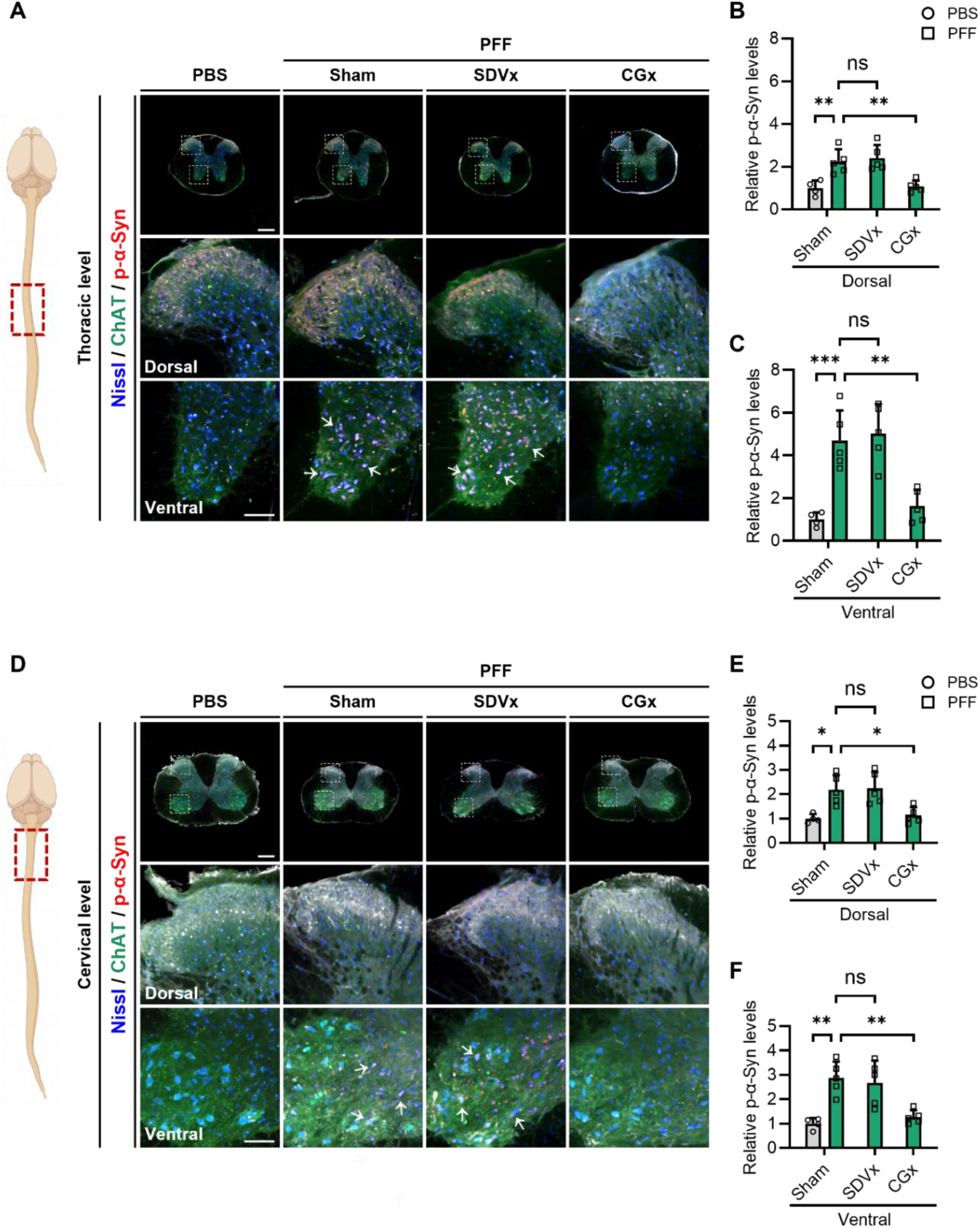
Celiac-superior mesenteric ganglionectomy prevents the accumulation and spread of α-synuclein aggregates in the spinal cord. **(A)** Schematic illustration of the mouse central nervous system indicating the thoracic spinal cord region of interest (red dashed box, left). Representative immunofluorescence images of coronal thoracic spinal cord sections collected at 10 months post-injection, stained for choline acetyltransferase (ChAT, green), p-α-Syn (red), and Nissl (blue) (right). High-magnification images of the dorsal horn (middle) and ventral horn (bottom) correspond to the white dashed boxes in the top panel. White arrows indicate p-α-Syn-positive inclusions in ChAT-positive motor neurons. Scale bars, 200 μm (top) and 50 μm (magnified). **(B and C)** Relative quantification of p-α-Syn levels in the dorsal horn (B) and ventral horn (C) of the thoracic spinal cord (*n* = 4 per group). **(D)** Schematic illustration of the mouse central nervous system indicating the cervical spinal cord region of interest (red dashed box, left). Representative immunofluorescence images of coronal cervical spinal cord sections collected at 10 months post-injection, stained for ChAT (green), p-α-Syn (red), and Nissl (blue) (right). High-magnification images of the dorsal horn (middle) and ventral horn (bottom) correspond to the white dashed boxes in the top panel. White arrows indicate p-α-Syn-positive inclusions in ChAT-positive motor neurons. Scale bars, 200 μm (top) and 50 μm (magnified). **(E and F)** Relative quantification of p-α-Syn levels in the dorsal horn (E) and ventral horn (F) of the cervical spinal cord (*n* = 4 per group). Data are presented as mean ± SEM. Statistical significance was determined by one-way ANOVA followed by Tukey’s multiple comparisons test; \**p* < 0.05, \*\**p* < 0.01, \*\*\**p* < 0.001; ns, not significant. PBS, phosphate-buffered saline; PFF, preformed fibrils; SDVx, subdiaphragmatic truncal vagotomy; CGx, celiac–superior mesenteric ganglionectomy; ChAT, choline acetyltransferase.

### Propagation of α-Syn PFF through both Parasympathetic and Sympathetic nerves

To directly track the neural conduits through which pathological α-Syn ascends from the gut, we evaluated the presence of p-α-Syn in peripheral neural structures 1 month following α-Syn PFF injection into the upper small intestine. We analyzed the Nodose Ganglion (NG), Vagus Nerve (VN), Celiac-Superior Mesenteric Ganglion (CSMG), and Splanchnic Nerve (SN) to assess propagation through both parasympathetic and sympathetic branches.

In the parasympathetic pathway, PFF injection induced a marked accumulation of p-α-Syn in HuC/D-positive neuronal cell bodies within the NG, as well as along Tuj1-positive nerve fibers in the trunk of the vagus nerve **(Figure 6A–6D)**. Quantitative analysis confirmed a significant increase in p-α-Syn signal intensity in both the NG and the vagus nerve in PFF-injected mice compared to PBS-injected controls **(Figure 6B and 6D)**.

**Figure 6.**
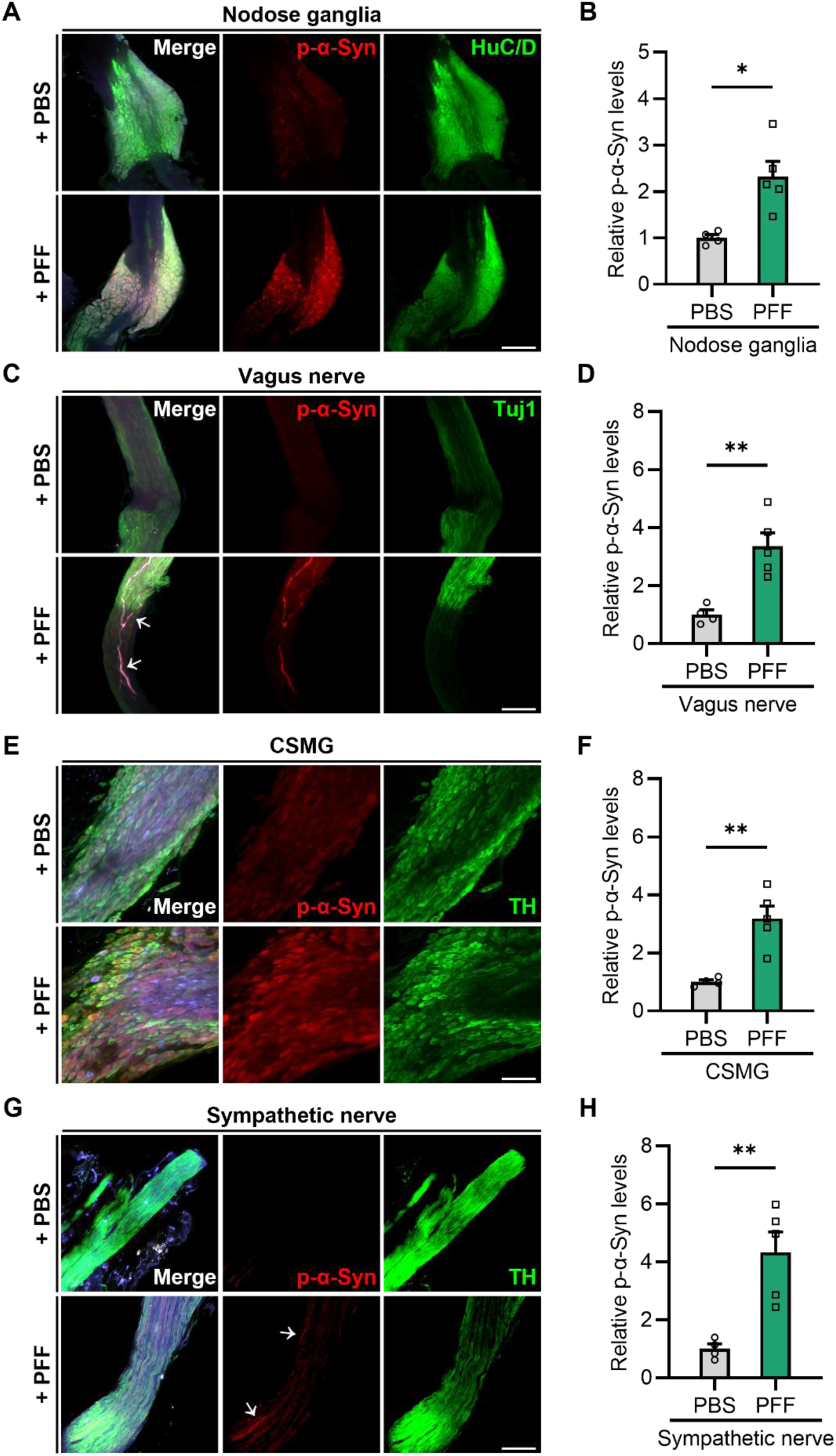
Gut-induced α-synuclein aggregates propagate along the vagus and spinal nerves. **(A)** Representative immunofluorescent images of whole-mounted nodose ganglia (NG) collected at 1 month post-injection following microinjection of *α*-synuclein preformed fibrils (*α*-Syn PFF) into the myenteric plexus of the duodenum, stained for HuC/D (green), p-α-Syn (red), and DAPI (blue). Scale bar, 100 *μ*m. **(B)** Relative quantification of p-α-Syn signal intensity in the nodose ganglion at 1 month post-injection (*n* = 4 per group). **(C)** Representative confocal immunofluorescent images of whole-mounted vagus nerve (VN) collected at 1 month post-injection following duodenal myenteric plexus injection of *α*-Syn PFF, stained for Tuj1 (green), p-α-Syn (red), and DAPI (blue). White arrows indicate p-α-Syn-positive pathology propagating along vagal nerve fibers. Scale bar, 100 *μ*m. **(D)** Relative quantification of p-α-Syn signal intensity in the vagus nerve at 1 month post-injection (*n* = 4 per group). **(E)** Representative confocal immunofluorescent images of whole-mounted celiac–superior mesenteric ganglia (CSMG) collected at 1 month post-injection following duodenal myenteric plexus injection of *α*-Syn PFF, stained for tyrosine hydroxylase (TH, green), p-α-Syn (red), and DAPI (blue). Scale bar, 100 *μ*m. **(F)** Relative quantification of p-α-Syn signal intensity in the CSMG at 1 month post-injection (*n* = 4 per group). **(G)** Representative confocal immunofluorescent images of whole-mounted sympathetic nerves collected at 1 month post-injection following duodenal myenteric plexus injection of *α*-Syn PFF, stained for TH (green), p-α-Syn (red), and DAPI (blue). White arrows indicate p-α-Syn-positive pathology propagating along sympathetic nerve fibers. Scale bar, 100 *μ*m. **(H)** Relative quantification of p-α-Syn signal intensity in sympathetic nerves at 1 month post-injection (*n* = 4 per group). Data are presented as mean ± SEM. Statistical significance was determined by unpaired two-tailed Student’s t-test; \**p* < 0.05, \*\**p* < 0.01, \*\*\**p* < 0.001; ns, not significant. PBS, phosphate-buffered saline; PFF, preformed fibrils; NG, nodose ganglion; VN, vagus nerve; CSMG, celiac–superior mesenteric ganglia; TH, tyrosine hydroxylase; Tuj1, neuron-specific class III beta-tubulin.

Similarly, examination of the sympathetic axis revealed prominent p-α-Syn pathology within the CSMG and splanchnic nerves **(Figure 6E–6H)**. In the CSMG, strong p-α-Syn signals co-localized with TH-positive sympathetic ganglionic neurons **(Figure 6E and 6F)**. Furthermore, p-α-Syn positive pathology was clearly observed extending along TH-positive axons in the splanchnic nerve, supporting active retrograde transport through sympathetic fibers **(Figure 6G and 6H)**.

Although our results did not reveal clear differences in propagation kinetics or route preference between the vagal/parasympathetic and spinal/sympathetic pathways, these observations at 1 month post-injection provide direct histological evidence that gut-administered α-Syn PFF propagates through both peripheral neural routes toward the central nervous system.

### Lewy bodies in the Spinal Cord of Parkinson’s Disease Patients

To determine whether the experimental observations in our animal models reflect pathological features relevant to human disease, and in particular whether pathological α-Syn aggregates are present in the spinal cord of PD patients, we examined human post-mortem CNS tissues, including the spinal cord **(Supplementary Table 1)**. Immunohistochemical analysis targeting p-α-Syn revealed a substantial burden of α-synucleinopathy in tissues derived from PD patients compared to age-matched healthy controls **(Figure 7A–7D)**. In the spinal cord of PD patients, prominent pS129-positive inclusions and aggregates reminiscent of Lewy bodies and Lewy neurites were detected **(Figure 7A)**. Quantitative assessment confirmed a significant elevation in the relative number of p-α-Syn-positive inclusions in the spinal cord of PD patients compared with healthy controls **(Figure 7B)**. Consistent with established PD neuropathology, dense p-α-Syn-positive pathology was also observed in the substantia nigra **(Figure 7C and 7D)**.

**Figure 7.**
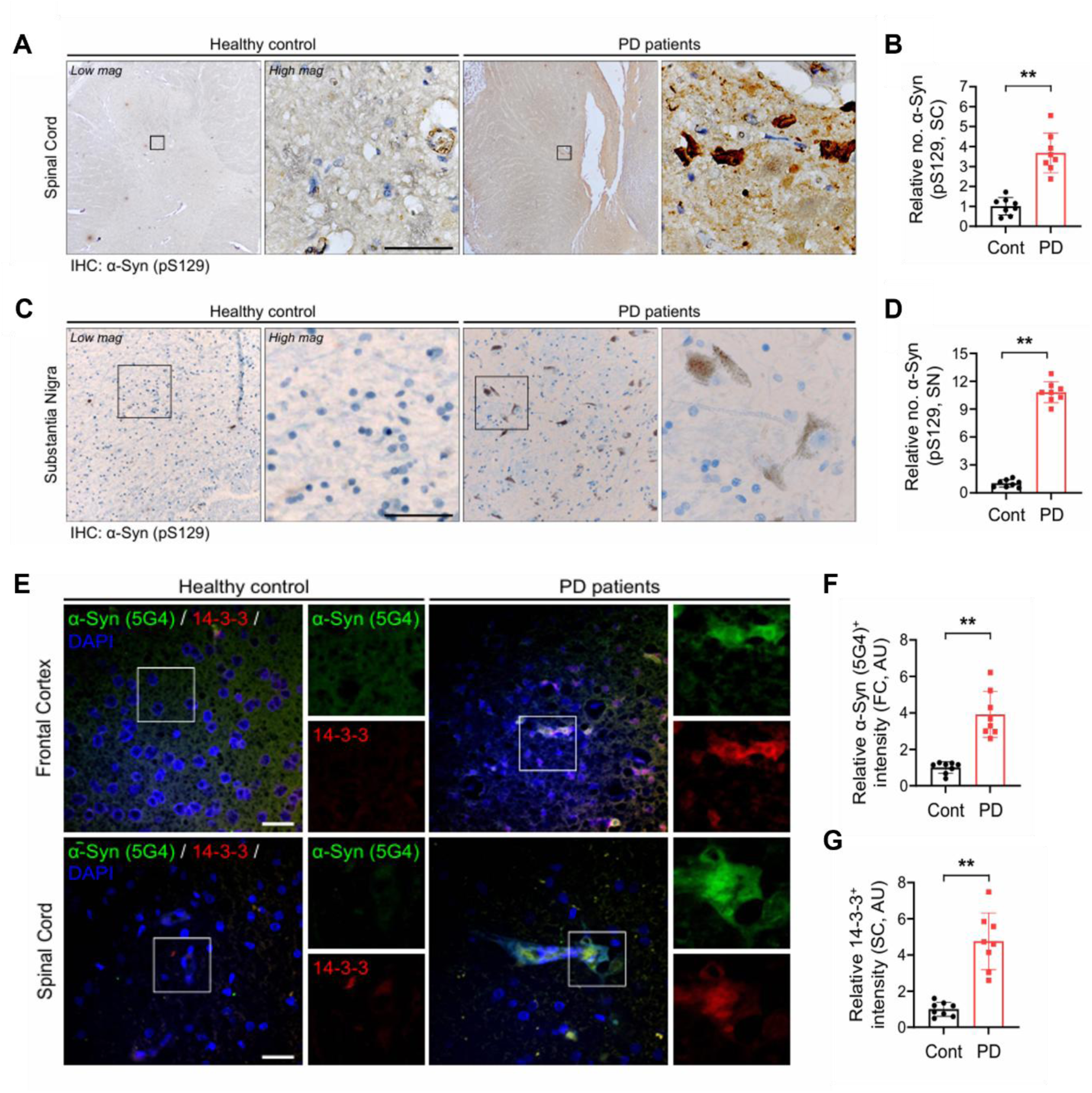
Pathological α-synuclein aggregates and Lewy body components accumulate in the spinal cord and brain of patients with Parkinson’s disease. **(A)** Representative DAB immunohistochemical (IHC) images of thoracic spinal cord sections from healthy controls and patients with Parkinson’s disease (PD), stained for p-α-Syn (pS129, brown) with hematoxylin counterstain (blue). Magnified images (high mag) correspond to the black dashed boxes in low-magnification panels (low mag). Scale bar, 50 *μ*m. **(B)** Relative quantification of p-α-Syn-positive inclusions in the spinal cord of healthy controls and PD patients (*n* = 8 cases per group). **(C)** Representative DAB IHC images of the substantia nigra (SN) from healthy controls and PD patients, stained for p-α-Syn (brown). High-magnification insets show Lewy body-like aggregates within dopaminergic area. Scale bar, 50 *μ*m. **(D)** Relative quantification of p-α-Syn-positive inclusions in the substantia nigra (*n* = 8 cases per group). **(E)** Representative confocal immunofluorescent images of human frontal cortex (top) and spinal cord (bottom) sections, co-stained for aggregated α-synuclein (5G4, green), 14-3-3 (red), and DAPI (blue). High-magnification panels show co-localization of 14-3-3 with 5G4-positive α-synuclein aggregates in PD patients. Scale bars, 25 *μ*m. **(F and G)** Relative signal intensity quantification of aggregated α-synuclein (5G4) in the frontal cortex (F) and 14-3-3 in the spinal cord (G) between healthy controls and PD patients (in arbitrary units, AU; *n* = 8 cases per group). Data are presented as mean ± SEM. Statistical significance was determined by unpaired two-tailed Student’s t-test; \**p* < 0.05, \*\**p* < 0.01, \*\*\**p* < 0.001; ns, not significant. Cont, healthy control; PD, Parkinson’s disease; SC, spinal cord; SN, substantia nigra; FC, frontal cortex; AU, arbitrary unit.

To further characterize the structural composition of these spinal cord inclusions, we performed immunofluorescence staining for misfolded α-Syn using the aggregate-specific antibody 5G4 together with 14-3-3, a well-established chaperone component of mature Lewy bodies **(Figure 7E–7G)**. In both the frontal cortex (FC) and the spinal cord of PD patients, strong 5G4-positive α-Syn aggregates co-localized with 14-3-3 immunoreactivity **(Figure 7E)**. Quantitative analysis showed a significant increase in 5G4-positive α-Syn aggregate intensity in the frontal cortex **(Figure 7F)** and a marked increase in 14-3-3 signal intensity in the spinal cord of PD patients compared to healthy controls **(Figure 7G)**. Together, these human post-mortem findings support that the presence of pathological α-Syn aggregates and mature Lewy body-like inclusions in the spinal cord of PD patients, consistent with the translational relevance of the spinal/sympathetic transmission pathway identified in our preclinical models.

## Discussion

Our study provides experimental evidence that gut-initiated α-synucleinopathy can access the CNS through anatomically parallel but context-dependent neural routes. Across two complementary mouse models, disruption of the prevertebral ganglia/splanchnic–spinal axis attenuated α-Syn pathology and associated behavioral deficits, although its relative contribution differed from that of the vagus across experimental contexts. The intracolonic rotenone model showed predominant dependence on the splanchnic–spinal axis, whereas gastroduodenal α-Syn PFF inoculation engaged both vagal and splanchnic–spinal pathways. Early detection of pathological α-Syn within peripheral structures associated with both routes further supported their parallel involvement. Together, these findings extend the prevailing vagus-centered model of body-first PD by identifying the splanchnic–spinal pathway as a complementary route to the CNS, and suggest that route-specific engagement may contribute to heterogeneity in autonomic patterns of Lewy pathology and in the biology of body-first PD.

The contribution of vagal and splanchnic-spinal pathways to gut-to-CNS α-syn pathology likely depends on multiple factors beyond connectivity alone, including disease subtype, regional gastrointestinal innervation, the mode of pathology initiation, and the molecular and cellular properties of the neurons encountered along each route. Body-first PD may itself include parasympathetic-and sympathetic-predominant trajectories **(Andersen et al., 2025)**, while vagal and spinal pathways show overlapping but regionally distinct innervation patterns across the gut **(Wang et al., 2020; Muller et al., 2020; Osman et al., 2023)**. PFF inoculation and rotenone exposure also initiate pathology through fundamentally different mechanisms **(Chmielarz and Domanskyi, 2021; Innos and Hickey, 2021)**. Moreover, successful propagation requires not only anatomical access but also uptake, intracellular persistence, axonal transport, release, and productive seeding in recipient neurons **(Freundt et al., 2012; Karpowicz et al., 2017)**. Thus, differential expression or activity of uptake and processing machinery, including LAG3, TLR2, and endolysosomal pathways, together with seed conformation, neuronal activity, and intrinsic susceptibility to α-syn seeding, may determine which peripheral route contributes most strongly to CNS α-synucleinopathy **(Mao et al., 2016; Kim et al., 2013, 2018; Emmenegger et al., 2021; Peelaerts et al., 2015; Yamada and Iwatsubo, 2018; Henderson et al., 2022; Courte et al., 2020; Luna et al., 2018)**.

How spinal α-Syn pathology subsequently engages the nigrostriatal system remains unresolved. Our denervation experiments establish a functional contribution of the splanchnic–spinal axis but do not define the longitudinal sequence of neuronal relays from the gut to the midbrain. Nevertheless, available neuroanatomy supports a distributed gut–spinal–brainstem network through which such propagation could occur. Polysynaptic tracing from the intestinal wall has identified gut-associated sympathetic circuits linking prevertebral ganglia with spinal and brainstem autonomic networks, including sympathetic premotor populations **(Muller et al., 2020)**. Complementing this gut-centered map, spinal-projecting neurons in the rostral ventromedial medulla directly innervate thoracic sympathetic preganglionic neurons and motor-related spinal regions **(Zhang et al., 2024)**. Defined neuronal populations in the nucleus tractus solitarius and medullary reticular formation likewise project to sympathetic preganglionic neurons in the thoracolumbar intermediolateral column **(Llewellyn-Smith et al., 2015)**, indicating that vagal-associated brainstem networks and spinal autonomic circuits are anatomically interconnected rather than fully segregated. More rostrally, the pedunculopontine and laterodorsal tegmental nuclei provide direct mesopontine input to midbrain dopaminergic neurons, including those in the substantia nigra pars compacta **(Gould et al., 1989; Futami et al., 1995; Xiao et al., 2016; French and Muthusamy, 2018)**. These connections raise the possibility that pathology entering thoracolumbar spinal circuits through splanchnic and prevertebral pathways could access interconnected medullary and pontine networks and ultimately reach the midbrain through polysynaptic relays. Further studies will be required to define whether and how pathological α-Syn propagates through this proposed gut–spinal–brain network.

Importantly, the translational relevance of this splanchnic–spinal conduit is powerfully underscored by our post-mortem human neuropathological analyses. While historical staging has focused on vagally connected lower brainstem structures, we observed a substantial burden of phosphorylated α-Syn inclusions within the thoracic spinal cord of PD patients **(Figure 7)**. Crucially, these spinal inclusions were not merely transient protein aggregates; rather, they exhibited robust co-localization between misfolded α-Syn (5G4) and 14-3-3, a definitive molecular chaperone signature of mature, consolidated Lewy bodies. The structural maturity of these spinal inclusions, mirroring the classical pathology observed in the substantia nigra and frontal cortex, confirms that the human spinal cord represents a major hub for stable synucleinopathy consolidation. The current clinical validation bridges our preclinical denervation findings with human disease, indicating that the splanchnic-spinal pathway is not a minor collateral route, but an active, pathophysiologically complete axis that underpins the systemic nature of body-first Parkinson’s disease.

## Limitation of Study

Several limitations should be considered. First, surgical denervation provides anatomical interruption rather than cell-type-specific manipulation. CGx disrupts multiple neural elements within the prevertebral ganglia/splanchnic axis, potentially affecting both sympathetic efferent and spinal afferent components, and therefore does not identify the neuronal population responsible for transmission. Second, SDVx and CGx alter gastrointestinal physiology and could influence α-Syn pathology indirectly through changes in motility, neural activity, inflammation, or the local tissue environment. Third, downstream pS129 α-Syn accumulation is consistent with propagation and seeding but does not directly trace the physical movement of individual α-Syn assemblies across successive relays. Fourth, the rotenone and PFF paradigms differ in both gastrointestinal site and nature of the initiating insult, precluding attribution of their differential pathway dependence solely to regional innervation. Finally, early peripheral mapping was performed at a single post-inoculation time point, and the available human material cannot establish the site of disease initiation or direction of spread. Longitudinal tracing combined with cell-type-specific pathway manipulation will be required to define the cellular routes and temporal sequence of gut-to-CNS α-Syn propagation.

## Materials and Methods

### Resource Availability

#### Lead contact

Further information and requests for resources and reagents should be directed to and will be fulfilled by the lead contact, Greg S. B. Suh.

#### Materials availability

This study did not generate new unique reagents. Materials and reagents used in this study are commercially available or available from the corresponding author upon reasonable request.

#### Data and code availability

All data supporting the findings of this study are available within the article and its supplemental information. Additional information required to reanalyze the data reported in this paper is available from the lead contact upon reasonable request. This study did not generate original code.

#### Experimental Model and Subject Details Animals

All animal experiments were conducted in accordance with institutional guidelines for the care and use of laboratory animals at the Korea Advanced Institute of Science and Technology (KAIST) and the Gwangju Institute of Science and Technology (GIST) and were approved by the respective Institutional Animal Care and Use Committees (IACUCs; KAIST protocol no. KA2024-091; GIST protocol no. GIST-2025-061). Male C57BL/6J mice (The Jackson Laboratory, stock no. 000664), 8–10 weeks of age at the start of experiments, were used for rotenone-and α-synuclein preformed fibril (PFF)-induced α-synucleinopathy studies. Mice were maintained under specific pathogen-free conditions and group-housed at 4–5 animals per cage in temperature-and humidity-controlled rooms (23 ± 1°C; 45–55% relative humidity) under a 12-h light/dark cycle (dark phase, 19:00–07:00). Standard chow (Teklad Global 18% Protein Rodent Diet, Envigo, 2018C) and water were provided ad libitum. Mice were acclimated to the animal facility for at least 3 days before experimental manipulation and were randomly assigned to experimental groups. Most experiments were performed at KAIST; the 1-month PFF tracing study used for peripheral pathway mapping was performed at GIST.

#### Human postmortem samples

Postmortem human central nervous system tissues from individuals with Parkinson’s disease (PD) and neurologically normal controls were obtained through the Emory University Parkinson’s disease research program. The cohort consisted of eight PD cases and eight controls, with demographic and clinical characteristics summarized in Supplementary Table 1. PD cases were neuropathologically confirmed by the presence of Lewy body pathology in the midbrain. Tissue samples analyzed in this study included the substantia nigra, frontal cortex, and thoracic spinal cord. Brain and spinal cord tissues were obtained from the same donor cohort. All tissue collection and research use were conducted under protocols approved by the relevant institutional ethics committees at Emory University.

### Method Details

#### Rotenone administration

Rotenone (Sigma-Aldrich, R8875) was freshly prepared on each day of administration in 4% carboxymethyl cellulose (CMC; Sigma-Aldrich, C5678) dissolved in 0.9% saline. The rotenone suspension was briefly homogenized and diluted with CMC vehicle to a final concentration of 7.5 mg/mL. Because rotenone gradually settled in suspension, the preparation was vortexed immediately before and periodically during administration to maintain uniformity. Mice received rotenone at 30 mg/kg once daily for 6 consecutive weeks, including weekends. Body weight was measured immediately before each daily administration, and the injection volume was adjusted accordingly (0.1 mL for a 25-g mouse; equivalent to 4 μL/g body weight). Vehicle-treated controls received an equivalent weight-adjusted volume of 4% CMC in 0.9% saline.

For intracolonic administration, mice were anesthetized with isoflurane (3–4% for induction and 2% for maintenance), and fecal material in the descending colon was gently cleared by abdominal massage before treatment. A PE-50 polyethylene tube (BD Intramedic, 427410; 8–9 cm in length) connected to a 23G needle and syringe was carefully inserted approximately 4–5 cm through the anus, reaching the anatomical bend of the descending colon (Figure S1A and S1B). The rotenone suspension was then administered slowly through the tubing. Following administration, the tail and posterior body were elevated for 5 min, positioning the anus upward to minimize leakage of the suspension before recovery from anesthesia. Vehicle-control mice underwent the identical anesthesia, tubing insertion, and post-administration procedure. Body weight was monitored daily throughout the 6-week treatment period and used to adjust the administered volume (Figure S1C).

For comparison with intracolonic delivery, a separate cohort received rotenone by oral gavage at the same dose (30 mg/kg), once daily for 6 consecutive weeks. Rotenone was prepared in the same 4% CMC vehicle at 7.5 mg/mL, and the administered volume was adjusted daily according to body weight. Rotenone was delivered directly into the stomach using a 20G oral gavage needle (JD-S-124-50-20G, 0.9 × 50 mm; Jeungdo Bio & Plant Co., Ltd.). Although anesthesia is not required for routine oral gavage, mice in this cohort were anesthetized with isoflurane using the same regimen as the intracolonic cohort to minimize procedural differences between the two administration paradigms. Vehicle-treated controls underwent the corresponding gavage procedure using an equivalent weight-adjusted volume of 4% CMC vehicle.

#### Generation of α-synuclein PFFs

Mouse α-synuclein preformed fibrils (PFFs) were generated and prepared as previously described (Kim et al., 2019). Briefly, recombinant mouse α-synuclein was fibrillized and subsequently sonicated to generate short fibrillar seeds suitable for neuronal uptake and propagation. PFFs were diluted in sterile phosphate-buffered saline (PBS) to a final concentration of 2.5 μg/μL immediately before gastrointestinal injection. The PFF preparation used in this study followed the same preparation and sonication conditions as those described previously **(Kim et al., 2019)**.

#### Gastroduodenal α-synuclein PFF injection

Gastrointestinal PFF injection was performed essentially as previously described (Kim et al., 2019). Mice were anesthetized with isoflurane and maintained on a heating pad during surgery. Following abdominal incision, the distal stomach and proximal duodenum were gently exposed. Sonicated mouse α-synuclein PFFs (2.5 μg/μL; total dose, 25 μg in 10 μL) were injected into four sites adjacent to the gastroduodenal myenteric plexus: two sites in the distal stomach and two sites in the proximal duodenum, approximately 0.5–1 cm from the pylorus. A volume of 2.5 μL was delivered at each site using a 10-μL Hamilton syringe, with each injection performed slowly over approximately 2.5 min. In the duodenum, the needle was inserted shallowly and approximately parallel to the longitudinal axis of the intestine to position the tip between the muscle layers while minimizing the risk of perforation. PBS-control mice received identical four-site injections of an equivalent volume of sterile PBS. Following injection, the abdominal wall and skin were closed, and mice were allowed to recover under postoperative monitoring.

#### Sham surgery

Mice were fasted for at least 3 h before surgery and anesthetized with isoflurane (3–4% for induction and 1.5–2% for maintenance). Abdominal hair was removed using depilatory cream, and the surgical area was disinfected with alcohol swabs. Mice were placed on a heating pad maintained at 36°C throughout the procedure. A midline skin incision of approximately 1.5 cm was made beginning ∼0.3 cm caudal to the xiphoid process, followed by a matching incision along the linea alba to open the abdominal wall. The skin and abdominal wall were gently retracted laterally using surgical retractors to expose the abdominal cavity.

For sham-operated mice, the abdomen was maintained open for approximately 30 min without transection or removal of neural structures. During this period, exposed abdominal organs were covered with sterile saline-moistened gauze and intermittently irrigated with sterile saline as needed to prevent tissue desiccation. The abdominal musculature was then closed with sutures, followed by closure of the skin. The surgical site was disinfected and treated with topical antimicrobial ointment. Mice were returned to their home cage with free access to food and water and were kept under an infrared warming lamp until fully recovered from anesthesia. Postoperative recovery was monitored by assessing spontaneous movement and general condition before the animals were returned to routine housing.

#### Subdiaphragmatic vagotomy

Bilateral subdiaphragmatic truncal vagotomy was performed using the same preoperative preparation, anesthesia, abdominal exposure, and postoperative closure procedures described for sham surgery. After opening the abdominal cavity, the liver lobes were gently displaced laterally using sterile saline-moistened cotton swabs to expose the stomach and distal esophagus. The stomach was carefully exteriorized and gently retracted to facilitate visualization of the esophagus, which was then followed proximally toward the diaphragm. Under a surgical microscope, the ventral vagal trunk was identified on the anterior surface of the esophagus using fine forceps and cotton swabs. The hepatic branch was identified as an anatomical landmark, and the main vagal trunk was isolated proximally to the hepatic branch. Approximately 1–2 mm of the vagal trunk was gently lifted away from the esophageal wall with fine forceps and transected using a cautery pen, while minimizing thermal injury to adjacent tissues. If the nerve was mechanically disrupted during isolation, both exposed nerve ends were cauterized to ensure complete interruption.

The esophagus was then gently rotated by approximately 90–120° using a saline-moistened cotton swab to expose the dorsal vagal trunk. The dorsal trunk was similarly identified, carefully separated from the esophageal surface proximal to the hepatic branch, and transected by cautery. After bilateral vagal transection, the operative field was inspected for bleeding or unintended tissue injury. When necessary, local hemostasis was achieved using hemostatic gauze. The stomach and liver were then returned to their anatomical positions, and the abdominal musculature and skin were closed as described for sham-operated mice. Postoperative care was performed according to the same protocol used for sham surgery.

To validate the completeness of vagotomy, after completion of the experimental paradigm, vagotomized mice received an intraperitoneal injection of the retrograde neural tracer FluoroGold (20 mg/kg in 0.9% saline; Fluorochrome). Successful vagotomy was confirmed by the absence of FluoroGold-labeled neurons in the dorsal motor nucleus of the vagus (DMV) (Figure S3A). As an additional physiological indicator of vagal disruption, gross stomach morphology and wet stomach weight were assessed at tissue collection, revealing the expected gastric enlargement following subdiaphragmatic vagotomy (Figure S4A and S4B).

#### Interruption of the prevertebral ganglia/splanchnic–spinal axis

Surgical interruption of the prevertebral ganglia/splanchnic–spinal axis was performed using the same preoperative preparation, anesthesia, abdominal exposure, and postoperative closure procedures described for sham surgery. After opening the abdominal cavity, the liver was gently displaced with sterile saline-moistened cotton swabs, and the small intestine was mobilized toward the animal’s right side to expose the left kidney and retroperitoneal vascular landmarks. When necessary, surgical retractors and saline-moistened gauze were used to maintain the viscera in this position. The left renal vein, superior mesenteric artery, celiac artery, and abdominal aorta were then identified under a surgical microscope to guide dissection of the prevertebral ganglia.

Using fine forceps, saline-moistened cotton swabs, and microsurgical scissors, connective and adipose tissues surrounding the left celiac ganglion were carefully removed to delineate the ganglion and its relationship to the adjacent vessels. The left celiac ganglion was then progressively dissected and removed from the periarterial tissue overlying the superior mesenteric artery. The dissection was extended along the anatomical continuity between the celiac and superior mesenteric ganglia, and as much of the superior mesenteric ganglion as could be safely accessed between the superior mesenteric artery and the left renal vein was removed. Complete excision of the superior mesenteric ganglion was not attempted when further dissection would have required manipulation beneath the superior mesenteric artery and posed a risk of major vascular injury.

The right celiac ganglion was subsequently identified within the region bounded by the celiac artery, superior mesenteric artery, and abdominal aorta and was similarly dissected and removed. After resection of the major prevertebral ganglia, fine periarterial neural fibers extending along the celiac artery, superior mesenteric artery, and abdominal aorta toward the pancreas and intestine were removed as extensively as safely possible. In addition, the major neural fibers extending from the left and right celiac ganglia toward the splanchnic nerves were identified and physically transected to interrupt continuity of the splanchnic pathway.

Following completion of the denervation procedure, the surgical field was inspected for bleeding and residual foreign material, and hemostasis was achieved as needed. Retractors and gauze were removed, the abdominal organs were returned to their anatomical positions, and the abdominal wall and skin were closed as described for sham-operated mice. Postoperative care was performed according to the same protocol used for sham surgery.

Successful disruption of the prevertebral ganglia was confirmed by gross anatomical inspection of the celiac–superior mesenteric ganglion region, with loss of the major ganglionic structures surrounding the celiac and superior mesenteric arteries (Figure S3B).

#### Pole test

Motor performance and bradykinesia were assessed using a vertical pole test. Mice were acclimated to the behavioral testing room for 30 min before testing. The apparatus consisted of a 50-cm vertical metal pole (1 mm in diameter) wrapped with gauze to provide sufficient grip. Before formal testing, mice were trained for two consecutive days, with three trials performed during each training session. A maximum cutoff time of 60 s was applied for each trial.

For the simplified pole-test paradigm used in Figure 2 and Figure S3, mice were placed near the top of the pole facing downward, and the time required to descend directly to the base of the pole was recorded as the total descent time. On the test day, each mouse underwent three trials, and the mean value was used for analysis.

For the conventional pole-test paradigm used in Figure 4, mice were placed approximately 5 cm below the top of the pole facing upward. The time required to orient downward was recorded as the turn time, and the subsequent time required to descend to the base of the pole was recorded as the descent time. Each mouse underwent three trials on the test day, and the mean values for turn time and descent time were used for statistical analysis.

#### Rotarod test

Motor coordination and balance were assessed using an accelerating rotarod test. Mice were trained on the rotarod apparatus for three consecutive days before testing. On the fourth day, mice were placed on the rotating rod, which accelerated from 4 to 40 rpm over 5 min, with a maximum trial duration of 5 min. The latency to fall from the rod was recorded. A trial was terminated when the mouse fell from the rod or when it passively rotated with the rod for two consecutive revolutions without attempting to maintain coordinated locomotion. Each mouse underwent two test trials, and the mean latency to fall was used for statistical analysis.

#### Grip strength test

Neuromuscular strength was assessed using a grip strength meter (BIO-GS3, Bioseb, FL, USA). Mice were allowed to grasp a metal grid with either the forelimbs alone or with both the forelimbs and hindlimbs. Once a stable grip was established, the mouse was gently pulled backward by the tail until the grip was released. The maximal force generated immediately before release was measured by the force transducer and recorded digitally in grams. Each mouse was tested three times, and the mean grip strength across the three trials was used for statistical analysis.

#### Spontaneous alternation Y-maze test

Spatial working memory was assessed using a spontaneous alternation Y-maze test as previously described (Kim et al., 2019). The maze consisted of three identical arms positioned at 120° angles, each measuring 40 cm in length, 10 cm in width, and 15 cm in wall height, and was constructed from opaque polyvinyl plastic. Each mouse was placed at the end of one arm and allowed to explore the maze freely for 8 min. The sequence and total number of arm entries were recorded manually. A spontaneous alternation was defined as consecutive entries into all three different arms (e.g., ABC, BCA, or CAB). The percentage of spontaneous alternation was calculated as:

Spontaneous alternation (%) = [number of alternations / (total arm entries − 2)] × 100.

The total number of arm entries was additionally used as an index of locomotor activity. The maze was cleaned with 10% ethanol between animals to minimize olfactory cues.

#### Nest building test

Nest-building behavior was assessed as an indicator of nigrostriatal sensorimotor function as previously described (**Fleming et al., 2004; Deacon, 2006**). A 2.5-g pressed cotton Nestlet was placed in the feeder of each cage. Retrieval of the nesting material required mice to rear and pull the cotton from the feeder using coordinated forelimb and orofacial movements, thereby engaging complex fine motor behavior. Nest-building performance was assessed 24 h after placement of the Nestlet. Nest quality was scored on a five-point scale based on the extent of Nestlet shredding and the organization and structure of the resulting nest, with higher scores indicating more complete nest construction.

#### Tail suspension test

The tail suspension test was performed with minor modifications from a previously described method (**Steru et al., 1985**). Mice were individually suspended by the tail inside an acrylic chamber (30 × 30 × 50 cm) using adhesive tape placed approximately 2 cm from the distal tip of the tail. The tape was attached to an overhead hook such that the mouse was suspended approximately 10 cm above the floor of the chamber. Each session was video-recorded for approximately 6 min and 10 s to ensure complete capture of the testing period. Immobility was manually scored during the first 6 min of the recording. Immobility was defined as the absence of active escape-directed movements; small body movements, passive tail rotation, and movements required for respiration were not scored as active movement. The cumulative immobility time during the 6-min test period was used for statistical analysis.

#### GI transit time assay

Whole-gut transit time was assessed using the carmine red assay as previously described, with minor modifications (Koester et al., 2021). Carmine red was prepared as a 6% (w/v) suspension in 0.5% methylcellulose and administered by oral gavage at 10 μL/g body weight (e.g., 250 μL for a 25-g mouse). Mice were fasted for 3 h before administration with free access to water, including a 30-min acclimation period in the behavioral testing room. Immediately after gavage, mice were individually placed in bedding-free cages. No observations were made during the first 60 min after administration; thereafter, cages were checked at 5-min intervals for the appearance of the first red-colored fecal pellet. Red coloration was initially assessed visually and, when necessary, confirmed by gently crushing the fecal pellet onto tissue paper. Gastrointestinal transit time was defined as the interval between carmine red administration and expulsion of the first confirmed red fecal pellet.

Fecal pellet output was assessed separately as an additional measure of gastrointestinal motility. Mice were individually transferred to cages containing a minimal amount of bedding sufficient to permit clear visualization and collection of fecal pellets and were acclimated for 30 min with ad libitum access to food and water. Following acclimation, fecal pellets were counted over a 1-h observation period. Cages were inspected approximately every 5–10 min, and newly excreted pellets were counted promptly to minimize loss due to coprophagy. The total number of fecal pellets produced during the 1-h observation period was used for statistical analysis.

#### Perfusion and tissue processing

Mice were deeply anesthetized with isoflurane until loss of reflexes, and adequate depth of anesthesia was confirmed by the absence of the pedal withdrawal reflex. Animals were then transferred to a surgical platform, where anesthesia was maintained by continuous isoflurane delivery. The limbs were secured with needles, and the thoracoabdominal surface was broadly disinfected with 70% ethanol. The thoracic cavity was opened by incision of the diaphragm and rib cage to expose the heart. Transcardial perfusion was first performed with ice-cold phosphate-buffered saline (PBS) containing heparin (10 U/mL).

For tissues intended for biochemical analyses, including immunoblotting and quantitative RT-PCR, perfusion was terminated after the PBS/heparin wash. Tissues, including colon, were rapidly dissected, snap-frozen in liquid nitrogen, and stored at −80°C until further processing. For histological analyses, PBS perfusion was followed by transcardial perfusion with 4% paraformaldehyde (PFA) in 1× PBS (pH 7.4). Brain, spinal cord, intestine, and other tissues of interest were then collected, rinsed in PBS, and post-fixed overnight in 4% PFA at 4°C before subsequent tissue-specific processing.

Peripheral neural tissues were dissected under a stereomicroscope using fine forceps and microsurgical scissors. Nodose ganglia were isolated with minor modifications from a previously described protocol for dissection of the vagus nerve and jugular–nodose ganglion complex (**Han and de Araujo, 2021**). Briefly, the occipital bone was split along the midline and the temporal bone was displaced from the occipital bone to expose the jugular–nodose ganglion complex and associated vagus nerve. The ganglion and adjacent vagal structures were then carefully dissected free from surrounding tissue. The cervical vagus nerve was additionally isolated by exposing the cervical region and tracing the nerve trunk alongside the esophagus and trachea toward its junction with the nodose ganglion.

The celiac–superior mesenteric ganglion (CSMG) complex and associated splanchnic nerve structures were isolated with minor modifications from a previously described protocol for isolation of the mouse sympathetic splanchnic–celiac–superior mesenteric ganglion complex (**Trevizan-Baú et al., 2024**). The CSMG region was first rapidly isolated using the celiac artery, superior mesenteric artery, abdominal aorta, and renal vasculature as anatomical landmarks. The dissected tissue was transferred to a PBS-filled Petri dish, where surrounding adipose, connective, and other non-neural tissues were carefully removed by fine microdissection. Neural structures extending from the bilateral prevertebral ganglionic region through the splanchnic pathway toward the spinal axis were subsequently isolated for downstream histological analysis.

#### Immunohistochemistry

Following transcardial perfusion and fixation as described above, brains were post-fixed in 4% paraformaldehyde for 16 h and cryoprotected in 30% sucrose in PBS (pH 7.4). Brains and Swiss-rolled colon tissues were embedded in OCT compound and sectioned at 30 μm using a cryostat. Free-floating sections were blocked in PBS containing 5% normal goat serum and 0.2% Triton X-100 and incubated with the appropriate primary antibodies. Brain sections were stained with an anti-tyrosine hydroxylase (TH) antibody (Abcam) to visualize midbrain dopaminergic neurons, whereas colon sections were pretreated with proteinase K before blocking to reduce proteinase K-sensitive α-synuclein species and enrich detection of proteinase K-resistant aggregates, followed by incubation with an anti-α-synuclein antibody (BD Biosciences).

Following primary antibody incubation, sections were incubated with the appropriate biotin-conjugated secondary antibody (anti-rabbit for TH staining and anti-mouse for α-synuclein staining). Immunoreactivity was visualized using SigmaFast DAB peroxidase substrate according to the manufacturer’s instructions. After mounting and imaging, TH-positive neurons in the midbrain were quantified using ImageJ.

#### Immunofluorescence

For colon immunofluorescence, Swiss-rolled colon tissues were embedded in OCT compound and sectioned at 30 μm using a cryostat. Free-floating sections were blocked in PBS containing 5% normal donkey serum and 0.2% Triton X-100 and subsequently incubated with primary antibodies overnight at 4°C in PBS containing 5% normal donkey serum. Colon sections were stained for α-synuclein or phosphorylated α-synuclein, together with DAPI nuclear counterstaining.

For spinal cord immunofluorescence, collected spinal cords were post-fixed in 4% paraformaldehyde for 16 h and cryoprotected in 30% sucrose in PBS (pH 7.4) for 48 h. Spinal cords were divided into cervical, thoracic, and lumbar regions, embedded in OCT compound, frozen, and sectioned at 30 μm using a cryostat. Brain tissues were processed similarly following post-fixation and cryoprotection, embedded in OCT compound, and serially sectioned at 30 μm. Free-floating brain and spinal cord sections were used for immunofluorescence staining, including analyses of the substantia nigra pars compacta (SNpc), locus coeruleus (LC), and dorsal motor nucleus of the vagus (DMV).

Following primary antibody incubation, sections were washed three times with PBS containing 0.1% Triton X-100 (PBST) and incubated with the appropriate fluorophore-conjugated secondary antibodies in PBS containing 5% normal donkey serum. Secondary antibody incubation was performed either overnight at 4°C or for 2 h at room temperature, depending on the staining condition. Alexa Fluor 488-, 555-, and 647-conjugated secondary antibodies were used as appropriate. Sections were then washed three times with 0.1% PBST, counterstained with DAPI, mounted using Dako fluorescent mounting medium, and imaged by confocal microscopy.

For whole-mount immunofluorescence of peripheral ganglia and nerve tissues used in Figure 6, isolated nodose ganglia, vagus nerves, celiac–superior mesenteric ganglion (CSMG) complexes, and associated splanchnic nerve tissues were post-fixed overnight in 4% paraformaldehyde and washed in PBS. Samples were permeabilized in 0.5% Triton X-100 in PBS (PBST) for 2 h at room temperature and subsequently blocked overnight at room temperature in 5% normal donkey serum prepared in 0.3% PBST. Primary antibodies were diluted in the same blocking buffer and incubated with the tissues for 48 h at 4°C. Samples were then washed three times in 0.1% PBST for 1 h per wash at room temperature. Appropriate fluorophore-conjugated secondary antibodies were diluted in blocking buffer and incubated with the tissues for an additional 48 h at 4°C, followed by three 1-h washes in 0.1% PBST at room temperature. Samples were counterstained with DAPI.

Following immunolabeling, tissues were cleared using a clearing-enhanced 3D (Ce3D) approach adapted from Li et al. (2019). Samples were incubated in Ce3D clearing solution for 24 h at room temperature, whole-mounted, and subsequently imaged by confocal microscopy.

#### Image acquisition and quantitative analysis

Immunofluorescence images were acquired using a Nikon AX Ti2 confocal microscope equipped with a 20× objective and NIS-Elements AR software. For sectioned tissues, Z-stack images spanning the full 30-μm tissue thickness were acquired at 5-μm intervals. Where necessary, tiled image acquisition was performed with 10% overlap between adjacent fields, followed by image stitching to reconstruct larger anatomical regions. Within each experiment, acquisition parameters, including laser power, detector gain, and offset, were kept constant across experimental groups. Maximum-intensity projections of the Z-stacks were generated for subsequent quantitative analysis.

Fluorescence intensity was quantified using NIS-Elements AR or ImageJ after background subtraction. Signal intensity was expressed as integrated density within defined regions of interest (ROIs). For brain regions, ROIs corresponding to the substantia nigra pars compacta (SNpc), locus coeruleus (LC), and dorsal motor nucleus of the vagus (DMV) were defined with reference to anatomical atlas landmarks. For spinal cord analyses, ROIs encompassing the dorsal or ventral horn were manually delineated based on the anatomical boundaries of each section. Cell counting was performed manually.

For whole-mount imaging of peripheral ganglia and nerves used in Figure 6, samples were imaged using the same Nikon AX Ti2 confocal microscope with a 20× objective. A region in which the ganglion or nerve structure was optimally visualized was selected as the central focal plane, and Z-stack images were acquired at 10-μm intervals over an approximately 140-μm depth, extending ∼70 μm above and below the central plane. Maximum-intensity projections were generated for visualization and quantitative analysis.

#### Tissue lysate preparation

Snap-frozen tissues stored at −80°C were lysed in ice-cold RIPA buffer containing 1% Triton X-100, 0.5% sodium deoxycholate, 0.1% SDS, 150 mM NaCl, and 50 mM Tris-HCl (pH 7.4), supplemented with protease and phosphatase inhibitor cocktails (Sigma-Aldrich). The volume of lysis buffer was adjusted according to tissue weight. Samples were thoroughly homogenized using a 150-W handheld homogenizer (Fisherbrand; product no. 15575809) and subsequently sonicated using a Q125 sonicator (Qsonica) at 20% amplitude for a total sonication time of 1 min using 5-s on/5-s off pulses. Lysates were centrifuged at 16,000 × g for 15 min at 4°C, and the resulting supernatants were collected. Protein concentrations were determined using a Pierce BCA Protein Assay Kit (Thermo Fisher Scientific). Lysates were then mixed with 4× SDS sample loading buffer containing reducing agent and adjusted to a final protein concentration of 2 μg/μL. Samples were heated at 75°C for 5 min and stored until subsequent immunoblot analysis.

#### Immunoblot analysis

Protein samples were resolved on laboratory-prepared 10% polyacrylamide gels. Twenty micrograms of total protein (10 μL of sample at 2 μg/μL) were loaded per lane and electrophoresed in 1× SDS running buffer using a Bio-Rad electrophoresis system at a constant current of 20 mA per gel for approximately 60 min. Proteins were subsequently transferred onto polyvinylidene difluoride (PVDF) membranes using a Bio-Rad wet-transfer system. Before transfer, PVDF membranes were activated in 100% methanol and equilibrated in distilled water. Wet transfer was performed in laboratory-prepared 1× transfer buffer containing 20% methanol at a constant voltage of 100 V for 90 min.

Following transfer, membranes were fixed in 4% paraformaldehyde (PFA) for 30 min at room temperature using a procedure adapted from a previously described membrane-fixation method (Sasaki et al., 2015). Membranes were then washed three times for 5 min each with distilled water and blocked for 1 h at room temperature in 5% skim milk prepared in Tris-buffered saline containing 0.1% Tween-20 (TBST). Primary antibodies were diluted in the same blocking solution and incubated with the membranes overnight at 4°C. Membranes were washed three times for 10 min each with TBST and incubated with the appropriate horseradish peroxidase (HRP)-conjugated secondary antibodies in blocking solution for 1 h at room temperature. After three additional 10-min washes with TBST, immunoreactive bands were detected using Pierce ECL Pico chemiluminescent substrate and imaged using a Bio-Rad ChemiDoc MP imaging system.

Band intensities were quantified using Bio-Rad Image Lab software or ImageJ. Integrated band intensity for each target protein was normalized to the corresponding β-actin signal, and normalized values were expressed as fold change relative to the appropriate control group.

#### RNA isolation and qRT-PCR

Total RNA was isolated from tissue samples using a standard TRIzol-based extraction method. Briefly, an appropriate volume of TRIzol reagent was added according to tissue weight, and samples were homogenized thoroughly. Chloroform was then added, samples were mixed vigorously, and phase separation was achieved by centrifugation at 13,000 × g for 15 min at 4°C. The aqueous phase was carefully collected, and RNA was precipitated using isopropanol, followed by ethanol washing. The resulting RNA pellet was resuspended in RNase-free water. RNA concentration and purity were assessed using a NanoDrop spectrophotometer.

Purified RNA was treated with DNase I to remove residual genomic DNA. One microgram of total RNA from each sample was then reverse-transcribed into complementary DNA (cDNA) using a Thermo Fisher Scientific cDNA synthesis kit according to the manufacturer’s instructions. Quantitative real-time PCR was performed using gene-specific primer pairs for Snca, Il6, and Gapdh. Relative gene expression was normalized to Gapdh and expressed as fold change relative to the appropriate control group.

#### Quantification and Statistical Analysis

All statistical analyses were performed using GraphPad Prism 11 (GraphPad Software). Data are presented as mean ± SEM unless otherwise indicated. Statistical significance between two groups was assessed using an unpaired two-tailed Student’s t-test. Comparisons among multiple groups were performed using one-way or two-way ANOVA followed by Tukey’s, Sidak’s, or Bonferroni multiple-comparisons test, as appropriate and as specified in the corresponding figure legends. A p value < 0.05 was considered statistically significant. Exact sample sizes and statistical tests are provided in the corresponding figure legends.

## Supporting information

Jung_et_al_supplement

## Acknowledgements

We thank Dong Woo Kim and Haeun Kang for their assistance with the rotenone experiments and for their contributions to the initial experimental setup. We acknowledge the KAIST behavioral core facility for technical support with mouse behavioral experiments. This work was supported by grants from the National Research Foundation of Korea (NRF) (NRF-2022M3A9F308298213, RS-2023-00255126, and RS-2024-00404132 to G.S.B.S.; RS-2023-00274696 to B.-K.J.) and by the Institute for Basic Science (IBS-R042-D1 to G.S.B.S.).

## Author contributions

G.S.B.S. conceived the project, designed experiments and interpreted the results across the study with B.-K.J. B.-K.J. conducted most experiments and analyzed data. M.K.S. and Y.-S.K. developed the intracolonic rotenone model. Y.K., H.P., and T.-I.K. designed and performed experiments using α-synuclein PFF. M.Y. performed imaging experiments. D.K.H. and S.S.K. performed the human postmortem studies. E.H.A. provided α-synuclein PFFs and fluorescent PFF. B.-K.J. and G.S.B.S. wrote the manuscript.

