## Supplementary material for "Splanchnic-spinal transmission of gut-initiated α-synucleinopathy": Jung_et_al_supplement

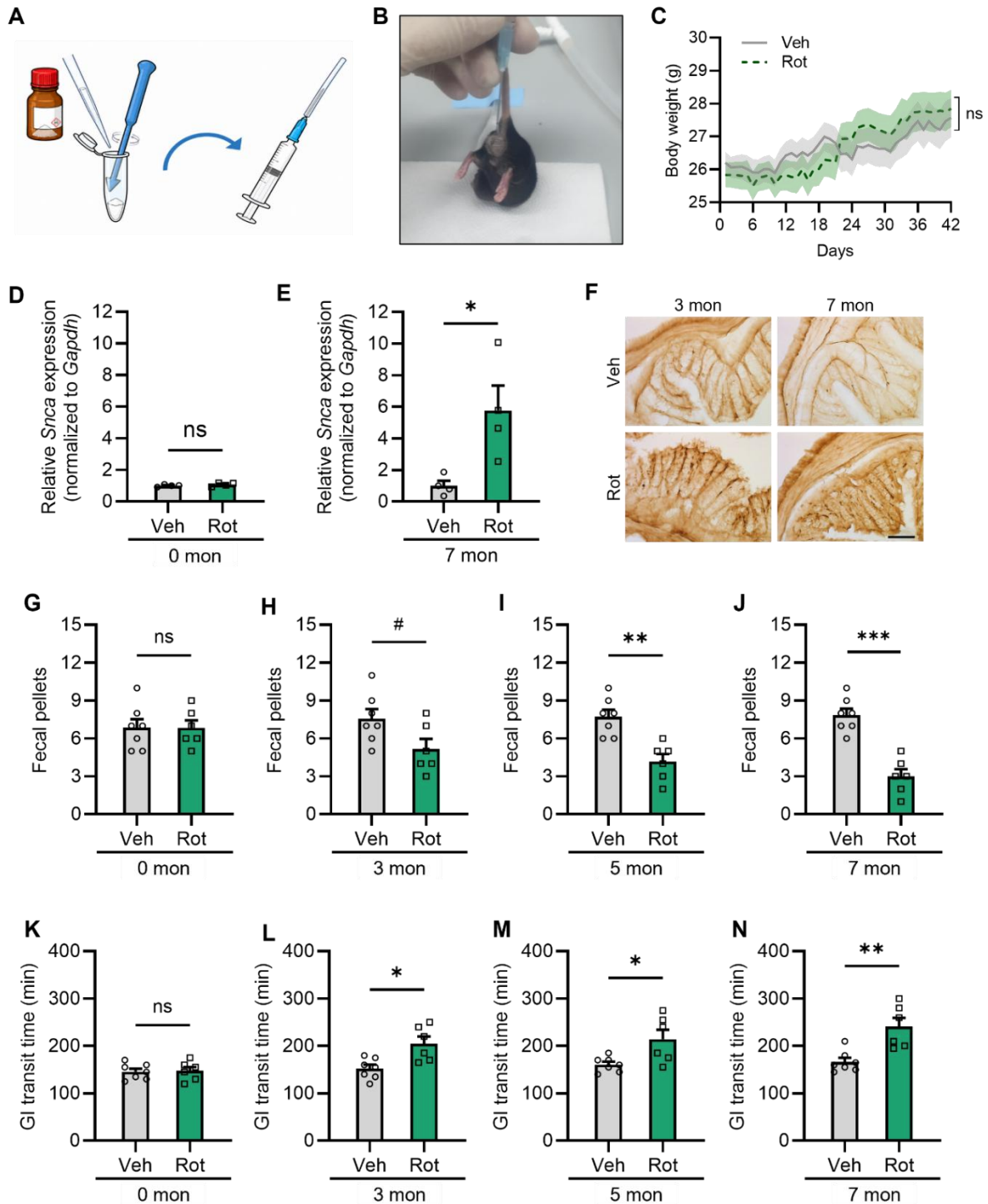

**Figure S1. Additional characterization of the intracolonic rotenone model, related to Figure 1.**

**(A and B)** Schematic illustration (A) and representative photograph (B) of the intracolonic rotenone administration procedure.

**(C)** Longitudinal body weight changes in vehicle (Veh)- and rotenone (Rot)-treated mice following intracolonic administration (n = 6 mice per group).

**(D and E)** Quantitative RT-PCR analysis of colonic *Snc*a mRNA expression in Veh- and Rot-treated mice at 0 mon (D) and 7 mon post-treatment (E) (n = 4 per group).

**(F)** Representative DAB immunohistochemical images of proteinase K (PK)-treated colon sections collected at 0 mon and 7 mon post-treatment to visualize PK-resistant α-Syn aggregates in Veh- and Rot-treated mice. Scale bar, 100 μm.

**(G–J)** Fecal pellet output measured over a fixed observation period in Veh- and Rot-treated mice at 0 mon (G), 3 mon (H), 5 mon (I), and 7 mon (J) post-treatment (n = 6–7 mice per group).

**(K–N)** Whole gastrointestinal (GI) transit time measured using the carmine red assay in Veh- and Rot-treated mice at 0 mon (K), 3 mon (L), 5 mon (M), and 7 mon (N) post-treatment (n = 6–7 mice per group).

Data are presented as mean ± SEM. Statistical significance was determined using an unpaired two-tailed Student's t-test or two-way ANOVA followed by Sidak's multiple-comparisons test, as appropriate; \*$p$ < 0.05, \*\*$p$ < 0.01, \*\*\*$p$ < 0.001; ns, not significant. Veh, vehicle; Rot, rotenone; PK, proteinase K; GI, gastrointestinal.

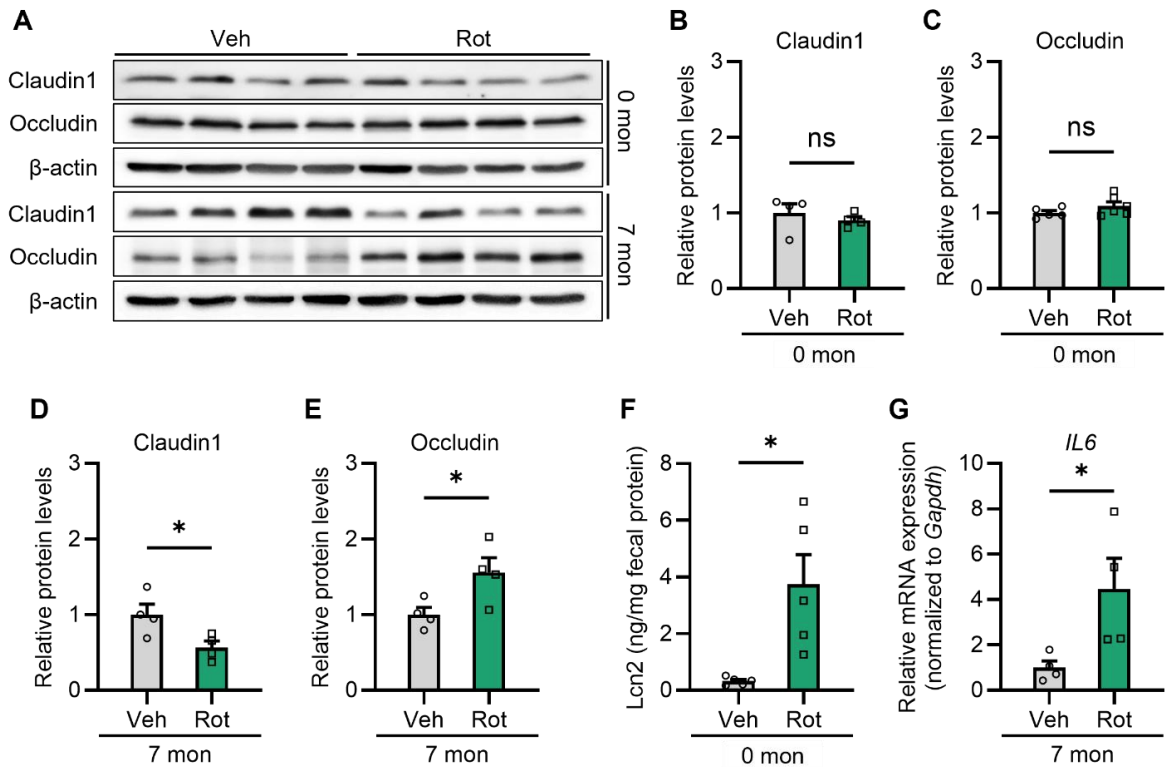

**Figure S2. Intracolonic rotenone administration alters intestinal epithelial junction markers and inflammatory responses, related to Figure 1.**

(A) Representative immunoblots of the tight junction proteins Claudin-1 and Occludin in colon tissues from vehicle (Veh)- and rotenone (Rot)-treated mice at 0 mon and 7 mon post-treatment. β-actin was used as a loading control.

(B and C) Quantification of colonic Claudin-1 (B) and Occludin (C) protein levels normalized to β-actin at 0 mon ($n = 4-6$ per group).

(D and E) Quantification of colonic Claudin-1 (D) and Occludin (E) protein levels normalized to β-actin at 7 mon post-treatment ($n = 4$ per group).

(F) ELISA quantification of fecal lipocalin-2 (Lcn2) levels, normalized to fecal protein, at 0 mon in Veh- and Rot-treated mice ($n = 5$ per group).

(G) Quantitative RT-PCR analysis of colonic *Il6* mRNA expression normalized to *Gapdh* at 7 mon post-treatment in Veh- and Rot-treated mice ($n = 4$ per group).

Data are presented as mean ± SEM. Statistical significance was determined using an unpaired two-tailed Student's t-test; $*p < 0.05$; ns, not significant. Veh, vehicle; Rot, rotenone; ELISA, enzyme-linked immunosorbent assay.

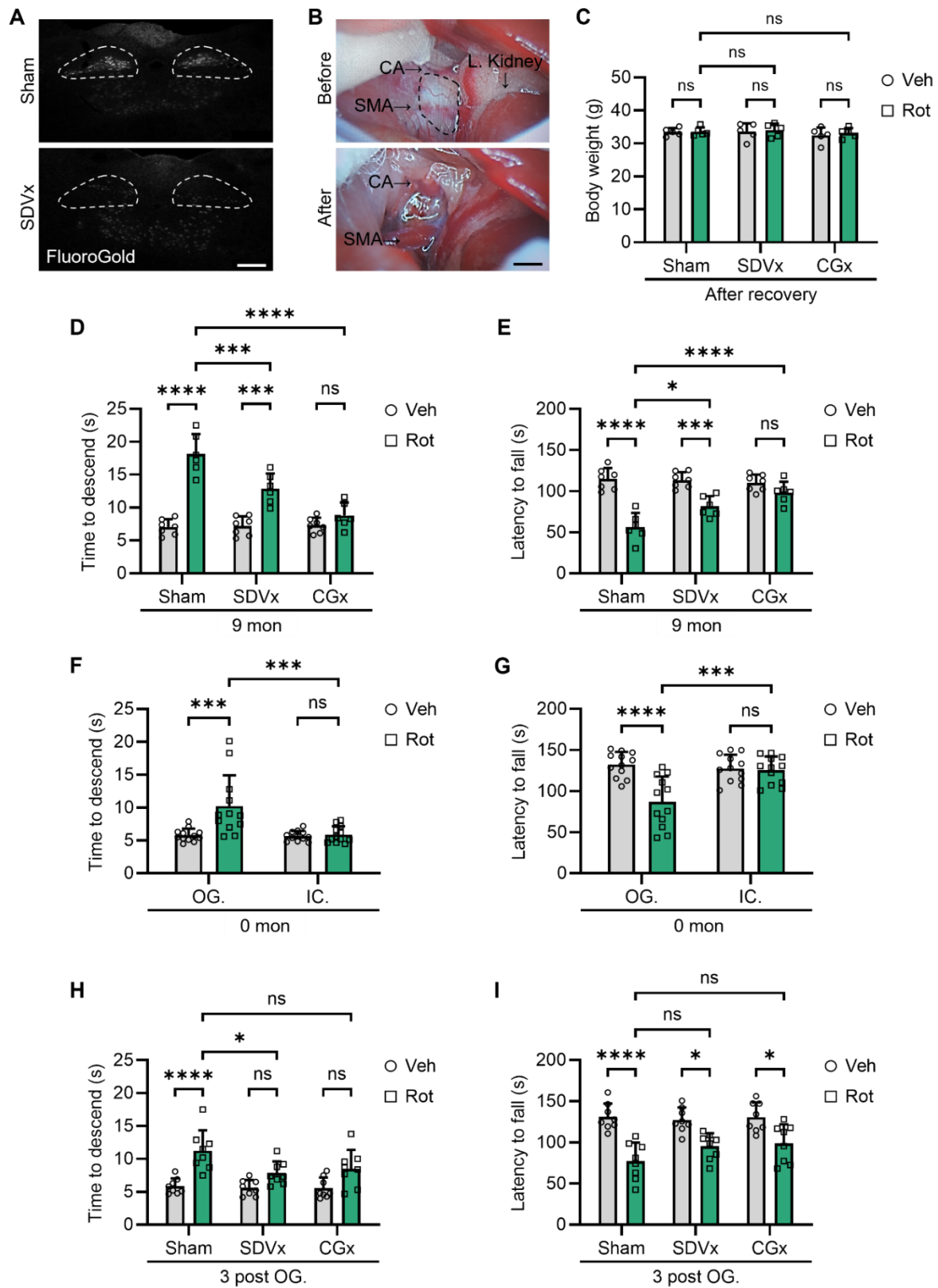

**Figure S3. Surgical validation of SDVx and CGx and behavioral evaluations following oral gavage or intracolonic rotenone administration, related to Figure 2.**

**(A)** Surgical validation of subdiaphragmatic truncal vagotomy (SDVx) by retrograde FluoroGold (FG) tracing in the dorsal motor nucleus of the vagus (DMV). Representative fluorescence images show retrogradely labeled neurons in Sham-operated mice (top) but not in SDVx-operated mice (bottom). Scale bar, 100 μm.

**(B)** Surgical validation of celiac–superior mesenteric ganglionectomy (CGx). Representative gross anatomical images show the intact celiac–superior mesenteric ganglion complex surrounding the celiac artery (CA) and superior mesenteric artery (SMA) before excision (top, dashed outline) and the surgical field after ganglion removal (bottom). Scale bar, 2 mm.

**(C)** Body weight recovery following Sham, SDVx, or CGx surgery as an index of postoperative physiological recovery (n = 5 mice per group).

**(D and E)** Pole-test (D) and rotarod (E) performance in vehicle (Veh)- and rotenone (Rot)-treated mice at 9 mon post-treatment following Sham, SDVx, or CGx surgery (n = 6–7 mice per group).

**(F and G)** Early behavioral comparison of oral gavage (OG.) and intracolonic (IC.) rotenone administration using the pole test (F) and rotarod test (G) at 0 mon (n = 12 mice per group).

**(H and I)** Pole-test (H) and rotarod (I) performance in rotenone-treated mice following Sham, SDVx, or CGx surgery at 3 mon after oral gavage administration (n = 8 mice per group).

Data are presented as mean ± SEM. Statistical significance was determined using one-way or two-way ANOVA followed by Tukey's multiple-comparisons test, as appropriate; \*$p$ < 0.05, \*\*$p$ < 0.01, \*\*\*$p$ < 0.001; ns, not significant. Veh, vehicle; Rot, rotenone; SDVx, subdiaphragmatic truncal vagotomy; CGx, celiac–superior mesenteric ganglionectomy; FG, FluoroGold; CA, celiac artery; SMA, superior mesenteric artery; OG., oral gavage; IC., intracolonic administration.

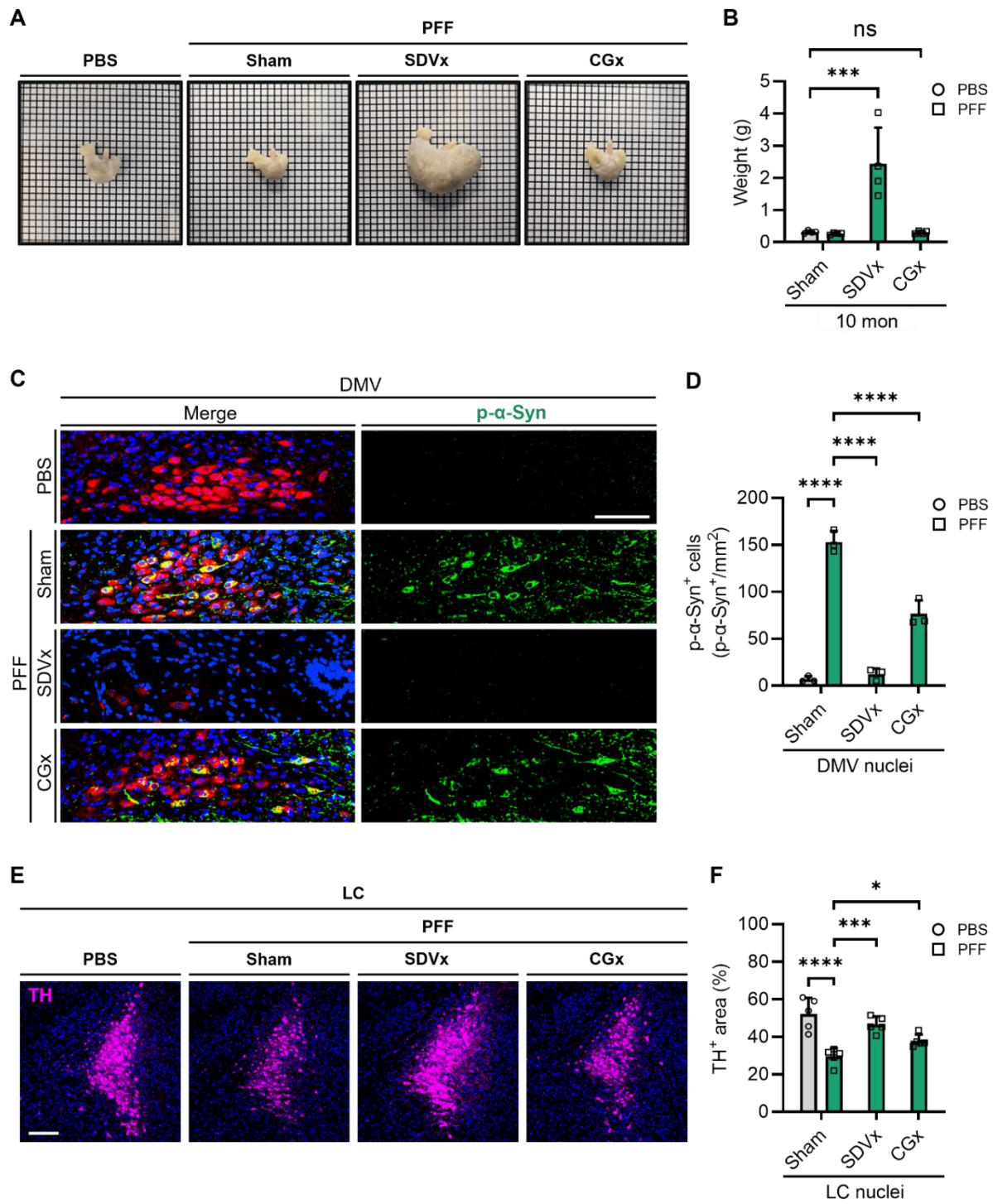

**Figure S4. Characterization of gastric phenotype and neuropathological evaluation in the brainstem following surgical denervation, related to Figure 4.**

**(A)** Representative gross anatomical photographs of stomachs isolated from mice in the phosphate-buffered saline (PBS), α-synuclein pre-formed fibrils (PFF), PFF with subdiaphragmatic truncal vagotomy (PFF + SDVx), and PFF with celiac–superior mesenteric ganglionectomy (PFF + CGx) groups. Grid scale, 1 mm per square.

**(B)** Quantitative measurement of wet stomach weight across experimental groups ($n = 4$ mice per group).

**(C)** Representative confocal immunofluorescent images of the dorsal motor nucleus of the vagus nerve (DMV) in the brainstem, stained for p-α-Syn (green), choline acetyltransferase (ChAT, red), and DAPI (blue). Scale bar, 100 μm.

**(D)** Relative quantification of p-α-Syn signal intensity in ChAT-positive cholinergic neurons within the DMV ($n = 3$ per group).

**(E)** Representative confocal immunofluorescent images of the locus coeruleus (LC) in the brainstem, stained for tyrosine hydroxylase (TH, magenta), and DAPI (blue). Scale bar, 100 μm.

**(F)** Relative quantification of p-α-Syn signal intensity in TH-positive noradrenergic neurons within the LC ($n = 4$ per group).

Data are presented as mean ± SEM. Statistical significance was determined by one-way ANOVA followed by Tukey's multiple comparisons test; \*$p < 0.05$, \*\*$p < 0.01$, \*\*\*$p < 0.001$; ns, not significant. PBS, phosphate-buffered saline; PFF, pre-formed fibrils; SDVx, subdiaphragmatic truncal vagotomy; CGx, celiac–superior mesenteric ganglionectomy; DMV, dorsal motor nucleus of the vagus nerve; LC, locus coeruleus; ChAT, choline acetyltransferase; TH, tyrosine hydroxylase.

| Sample | Age at onset | Age at death | Duration | ApoE allele | Sex |
| --- | --- | --- | --- | --- | --- |
| Control 1 |  | 91 |  | E3/3 | Female |
| Control 2 |  | 70 |  | E3/3 | Male |
| Control 3 |  | 72 |  | E3/3 | Male |
| Control 4 |  | 69 |  | E3/3 | Male |
| Control 5 |  | 84 |  | E2/3 | Female |
| Control 6 |  | 65 |  | E3/3 | Female |
| Control 7 |  | 75 |  | E3/3 | Female |
| Control 8 |  | 59 |  | E2/3 | Male |
| PD 1 | 61 | 72 | 11 | E3/4 | Female |
| PD 2 | 77 | 90 | 13 | E3/3 | Female |
| PD 3 | 54 | 75 | 21 | E2/3 | Male |
| PD 4 | 48 | 69 | 21 | E3/3 | Male |
| PD 5 | 64 | 84 | 20 | E3/3 | Male |
| PD 6 | 61 | 75 | 14 | E2/3 | Female |
| PD 7 | 52 | 58 | 6 | E4/4 | Male |
| PD 8 | 37 | 77 | 40 |  | Female |

PD, Parkinson's disease; ApoE, apolipoprotein E; N/A, not available.

*(Note: Age at onset, age at death, and disease duration are expressed in years.)*

**Supplementary Table 1. Demographic and clinical characteristics of human brain tissue donors, related to Figure 7.**
